# A guanidine-based coronavirus replication inhibitor which targets the nsp15 endoribonuclease and selects for interferon-susceptible mutant viruses

**DOI:** 10.1101/2024.09.09.611961

**Authors:** Benjamin Van Loy, Eugènia Pujol, Kenichi Kamata, Xiao Yin Lee, Nikolai Bakirtzoglou, Ria Van Berwaer, Julie Vandeput, Cato Mestdagh, Leentje Persoons, Brent De Wijngaert, Quinten Goovaerts, Sam Noppen, Maarten Jacquemyn, Kourosh Ahmadzadeh, Eline Bernaerts, Juan Martín-López, Celia Escriche, Bert Vanmechelen, Besir Krasniqi, Abhimanyu K. Singh, Dirk Daelemans, Piet Maes, Patrick Matthys, Wim Dehaen, Jef Rozenski, Kalyan Das, Arnout Voet, Santiago Vázquez, Lieve Naesens, Annelies Stevaert

## Abstract

The approval of COVID-19 vaccines and antiviral drugs has been crucial to end the global health crisis caused by SARS-CoV-2. However, to prepare for future outbreaks from drug-resistant variants and novel zoonotic coronaviruses (CoVs), additional therapeutics with a distinct antiviral mechanism are needed. Here, we report a novel guanidine-substituted diphenylurea compound that suppresses CoV replication by interfering with the uridine-specific endoribonuclease (EndoU) activity of the viral non-structural protein-15 (nsp15). This compound, designated EPB-113, exhibits strong and selective cell culture activity against human coronavirus 229E (HCoV-229E) and also suppresses the replication of SARS-CoV-2. Viruses, selected under EPB-113 pressure, carried resistance sites at or near the catalytic His250 residue of the nsp15-EndoU domain. Although the best-known function of EndoU is to avoid induction of type I interferon (IFN-I) by lowering the levels of viral dsRNA, EPB-113 was found to mainly act via an IFN-independent mechanism, situated during viral RNA synthesis. Using a combination of biophysical and enzymatic assays with the recombinant nsp15 proteins from HCoV-229E and SARS-CoV-2, we discovered that EPB-113 enhances the EndoU cleavage activity of hexameric nsp15, while reducing its thermal stability. This mechanism explains why the virus escapes EPB-113 by acquiring catalytic site mutations which impair compound binding to nsp15 and abolish the EndoU activity. Since the EPB-113-resistant mutant viruses induce high levels of IFN-I and its effectors, they proved unable to replicate in human macrophages and were readily outcompeted by the wild-type virus upon co-infection of human fibroblast cells. Our findings suggest that antiviral targeting of nsp15 can be achieved with a molecule that induces a conformational change in this protein, resulting in higher EndoU activity and impairment of viral RNA synthesis. Based on the appealing mechanism and resistance profile of EPB-113, we conclude that nsp15 is a challenging but highly relevant drug target.

**Author summary:** Despite major achievements in ending the global COVID-19 crisis through vaccines and antiviral drugs, SARS-CoV-2 infection remains a serious health threat for vulnerable individuals, such as elderly and immunocompromised patients. These populations could benefit from innovative treatments that target the virus from multiple angles. In this study, we focus on the nsp15 endoribonuclease (EndoU), a poorly understood coronavirus protein that aids in viral evasion from the host innate immune response and is also assumed to regulate viral RNA synthesis. Our research centers around a newly identified, first-in-class inhibitor of coronavirus nsp15, designated EPB-113. Although nsp15 is best-known for its function in interferon evasion, EPB-113 mainly acts via an interferon-independent mechanism that is situated during viral RNA synthesis. When incubated with nsp15 protein, EPB-113 induces a structural change and enhances its EndoU enzyme activity. EPB-113 selects for escape mutants that carry substitutions in the core of EndoU, leading to a severe replication defect. Hence, our study introduces a novel drug concept to disturb the multifunctional nsp15 protein and suppress coronavirus replication on multiple fronts.

## Introduction

In the five years since COVID-19 was first identified, Severe Acute Respiratory Syndrome Coronavirus 2 (SARS-CoV-2) has caused an estimated 777 million human infections and 7.1 million deaths worldwide [1]. The fast roll-out of innovative vaccines was crucial in addressing the global health crisis when SARS-CoV-2 suddenly emerged in the human population. Today, annual COVID-19 vaccination remains recommended, as the vaccines provide only short-term immunity and the virus continues to undergo antigenic drift. Serious complications from COVID-19 are common in vulnerable individuals, such as the elderly or immunocompromised patients. For those with mild-to-moderate COVID-19 who are at high risk for progression to severe disease, antiviral medication is recommended, specifically ritonavir-boosted nirmatrelvir [which targets the coronavirus main protease (M^pro^)] or remdesivir [a nucleotide inhibitor of the viral RNA-dependent RNA polymerase (RdRp)] [2].

To prepare for future outbreaks caused by drug-resistant variants or novel zoonotic coronaviruses (CoVs), exploring alternative antiviral drug targets is essential. Many ongoing efforts are focused on one of the 16 non-structural proteins (nsps) that make up the viral RNA transcription-replication complex (RTC) [3–5]. Several nsps help the virus to evade the host innate immune response that aims to halt viral replication [6]. This dual function also applies to nsp15, a uridine-specific endoribonuclease (EndoU) that is highly conserved among all CoVs [7, 8]. As part of the RTC, nsp15 is believed to cleave specific sites in viral RNA to regulate viral RNA synthesis [9]. Additionally, EndoU prevents the accumulation of viral dsRNA intermediates, thereby avoiding their detection by dsRNA sensors of the innate immune response [10–12]. This may be accomplished by EndoU cleavage of the 5′-poly(U) extension in viral negative-sense RNA [12] or the CACA sequence near the 3′-poly(A) tail of the positive strand [9]. Evidence also suggests that nsp15 interacts with cellular proteins involved in the production of type I interferon (IFN-I) [13, 14] and may protect infected cells against apoptosis [10, 15], autophagy [16], cell cycle arrest [17] or the formation of stress granules [18–20].

The immune-evading activity of EndoU has been demonstrated in cell culture studies with different CoVs [10, 11, 21, 22], including human nasal epithelial cells infected with SARS-CoV-2 [23]. EndoU-deficient mutants of mouse hepatitis virus (MHV) and porcine epidemic diarrhea virus (PEDV) proved significantly less pathogenic than their wild-type (WT) counterparts in mice and piglets, respectively [10, 21]. However, an EndoU-deficient mutant of transmissible gastroenteritis virus exhibited comparable virulence to WT in piglets, while the analogous mutant of feline infectious peritonitis virus was more virulent than WT in cats [24]. In hamster models, an EndoU-deficient form (i.e. H234A_nsp15_-mutant) of SARS-CoV-2 generated lower viral titers in the airways, while showing similar pathology as or higher innate immune responses than the WT [25, 26]. Hence, many questions remain about the functions of EndoU in viral replication and immune evasion.

The potential of nsp15 as a novel drug target has spurred recent efforts to identify inhibitors in enzymatic EndoU assays [27–33] and resolve the protein structure of nsp15 in complex with a mono- or oligonucleotide ligand [34–38]. For nearly all CoVs, including SARS-CoV-2, nsp15 adopts a characteristic hexameric structure consisting of a dimer of trimers, with the six EndoU domains facing outwards of the complex [39]. Each monomer contains an N-terminal domain (NTD), middle domain and C-terminal domain harboring the EndoU site. Nsp15 hexamerization is regulated by the NTD and crucial for EndoU enzymatic activity [40]. A recent review covers the biochemical properties, structure and assumed functions of nsp15 in greater detail and can be found elsewhere [41].

Our interest in developing nsp15 inhibitors started several years ago, following the identification of a betulonic acid-based inhibitor (encoded 5h in reference [42]) that exhibits submicromolar activity against human coronavirus 229E (HCoV-229E), low cytotoxicity and a mechanism related to nsp15. The finding that viral resistance to this compound was mapped to an inter-protomer region in the hexameric nsp15 protein of HCoV-229E, allowed us to create a docking model and predict the binding mode of 5h [42]. Shortly after, we identified compound EPB-113, another inhibitor of HCoV-229E that is structurally unrelated to 5h. Intrigued by the discovery that viral resistance to EPB-113 was linked to mutations in the catalytic center of nsp15 EndoU, we conducted the present study to uncover the mechanism of this antiviral molecule. Using IFN responsive and non-responsive cell types as well as one-cycle replication assays, we discovered that our inhibitor mainly acts in an IFN-independent manner, with IFN boosting being an only minor effect. Biochemical assays revealed that EPB-113 directly binds to hexameric nsp15 protein, thereby enhancing its cleavage activity. Through thorough profiling of the EPB-113-resistant nsp15-mutant viruses, we explored the association between EndoU, the innate immune response and viral replication kinetics. Taken together, our findings suggest that EPB-113 acts through a previously unrecognized pharmacological mechanism to target nsp15 and enhance its EndoU activity up to a level that is not compatible with viral replication.

## Materials and methods

### Compounds

In S1 Appendix, we provide all details on the chemical synthesis, analysis and full characterization of diphenylurea compounds EPB-113, EPB-102 and JMLV-061. The synthesis of 5h, a closely related 1,2,3-triazolo-fused betulonic acid derivatives, was described previously [42].

The following compounds were purchased from a commercial supplier: GS-441524 (Carbosynth); E64d (MedChemExpress); nirmatrelvir (Chemat); and K22 (ChemDiv). Recombinant human IFN-β was purchased from PeproTech.

### Cells and media

Human embryonic lung fibroblast cells [HEL299; American Type Culture Collection (ATCC) No. CCL-137] were cultured in Dulbecco’s modified Eagle medium (DMEM) supplemented with 10% fetal calf serum (FCS); 10 mM HEPES; 1 mM sodium pyruvate; 6 mM L-glutamine; 0.1 mM non-essential amino acids (NEAA); and 25 mM glucose. For experiments, they were seeded in 96-well plates and grown until confluent. The infection medium was DMEM with 2% FCS; 1 mM sodium pyruvate; 0.075% sodium bicarbonate; 0.1 mM NEAA; 4 mM L-glutamine; 25 mM glucose; and pen-strep (= 100 U/ml penicillin and 0.1 mg/ml streptomycin). Human bronchial epithelial 16HBE cells (kind gift from P. Hoet, KU Leuven) and BHK-21J cells (donated by P. J. Bredenbeek) were maintained in DMEM with 10% FCS; 1 mM sodium pyruvate; 0.075% sodium bicarbonate; 0.1 mM NEAA; 4 mM L-glutamine; and 25 mM glucose. For experiments, 16HBE cells were seeded on day -1 in 96-well plates (30,000 cells per well), using the same medium but with 2% FCS and pen-strep. A549 cells expressing ACE2 and TMPRSS2 (A549^ACE2/TMPRSS2^) were purchased from InvivoGen (No. a549d-cov2r) and cultured in DMEM supplemented with 6 mM L-glutamine; 25 mM glucose; 10% FCS; 100 µg/ml hygromycin and 0.5 µg/ml puromycin. HEK-Blue IFN-α/β cells (kindly donated by X. Saelens, UGent) were grown in DMEM with 4 mM L-glutamine; 25 mM glucose; 10% heat-inactivated FCS; pen-strep; 30 µg/ml blasticidin; and 100 µg/ml zeocin.

Human macrophages were purified from healthy donor buffy coats, obtained from the Red Cross Belgium according to KU Leuven agreement RKOV_19006. After isolating the mononuclear cell fraction by density gradient centrifugation on Pancoll (PAN-Biotech), the monocytes were purified by adherence for 1.5 h at 37 °C, followed by three PBS washes to remove non-adherent cells. The cells were seeded in 96-well plates (90,000 cells per well) and allowed to differentiate into macrophages during six days incubation in RPMI1640 medium with 2 mM glutamine and supplemented with 10% heat-inactivated FCS; 2 mM L-alanyl-glutamine; 20 ng/ml macrophage-colony stimulating factor (M-CSF, PeproTech); and pen-strep. Fresh medium was provided on day 3. For infection experiments, we used the same medium but with 2% FCS.

The Huh7-STAT1-KO cell line was made by CRISPR/Cas9-mediated knockout in the Huh7 cell line, using a STAT1 sgRNA (5′-AGAACACGAGACCAATGGTG-3′) from the Human Brunello CRISPR knockout pooled library [43]. The guide sequence was cloned into the pLentiCRISPRv2 plasmid (Addgene, 52961), previously adapted to include a Pro-Code sequence [44]. HEK293T cells were seeded in 75-cm² culture flasks at 45% confluency and, on the next day, co-transfected using Turbofectin 8.0 (OriGene) with the pLentiCRISPR plasmid and the lentiviral packaging plasmids pMD2.G and psPAX2, to generate lentiviral particles coated with the VSV-G protein. Twenty-four hours post-transfection, the medium was changed to FCS-free medium supplemented with 1% BSA and 20 µg/ml gentamicin. The supernatants containing lentiviral particles were harvested 72 h post-transfection and stored at -80 °C. Huh7 cells were transduced with these STAT1 sgRNA-expressing lentiviruses, then selected with puromycin for three days. After the successful KO was verified by automated western blot (S3 Appendix and S1 Fig), the cells were cultured in DMEM-based medium supplemented with 10% FCS. For experiments, the cells were seeded on day -1 in 96-well plates (30,000 cells/well), using the same medium but with 5% FCS and pen-strep.

All cells were incubated in a humidified 5% CO_2_ incubator at 37 °C.

### Viruses

HCoV-229E (ATCC No. VR-740) was expanded in HEL299 cells. Reverse-engineered HCoV-229E virus carrying substitution H250A_nsp15_ was kindly provided by V. Thiel (University of Bern). The SARS-CoV-2 viruses were created by reverse-engineering (see below). The stocks of wild-type and nsp15-mutant viruses (see below) were titrated in HEL299 (for HCoV-229E) or A549^ACE2/TMPRSS2^ cells (for SARS-CoV-2) and the 50% cell culture infective dose (CCID_50_) was calculated by the method of Reed and Muench. Unless stated differently in the text, all infection experiments were done at 35 °C for HCoV-229E and 37 °C for SARS-CoV-2.

### Antiviral and cytotoxicity assays

Unless specified differently, the compounds’ antiviral activity was determined in HEL299 cells infected with HCoV-229E or in A549^ACE2/TMPRSS2^ cells infected with SARS-CoV-2. The compounds were serially diluted in medium, then added to semiconfluent cells in 96-well plates, immediately followed by the virus at a multiplicity of infection (MOI) of 100 CCID_50_. A mock-infected plate was prepared in parallel, to assess compound cytotoxicity. To study combinations, different concentrations of the two compounds were added in a matrix of 8×8 wells. For the CPE reduction assay, the viral CPE was scored by microscopy at day 5 p.i., then quantified by the colorimetric MTS cell viability assay. After replacing the supernatant by CellTiter 96AQ_ueous_ MTS Reagent (Promega) and incubating the plates for 24 h (HEL299) or 6 h (A549^ACE2/TMPRSS2^) at 37 °C, the A_490_ values were measured in a Tecan Spark 10M plate reader. To calculate the EC_50_ (50% antivirally effective concentration) and CC_50_ (50% cytotoxic concentration) values, we used the formulas specified elsewhere [45].

For virus yield reduction assays, the supernatants were harvested at day 3 to 5 p.i. To quantify the viral load by RT-qPCR [33], the samples were mixed with Cells-to-cDNA II lysis buffer (Invitrogen) and incubated for 10 min at 75 °C. The number of viral RNA genomes (= viral load) was determined by one-step RT-qPCR (iTaq Universal Probes One-Step kit, BioRad), employing N-gene specific primers and probe (see sequences provided in S2 Appendix) and an N-gene plasmid standard to allow absolute quantification [46, 47]. The RT-qPCR program consisted of: 10 min at 50 °C; 2 min at 95 °C; and 40 cycles of 15 s at 95 °C and 60 s at 60 °C. To quantify the yield of infectious virus, we incubated serial dilutions of the supernatants with HEL299 cells in 96-well plates, scored the CPE at day 5 p.i. and calculated the CCID_50_ titers as described above. Antiviral activity in these virus yield assays was expressed as the EC_90_ value, i.e. compound concentration producing 1-log_10_ reduction in the number of viral RNA genomes or in the CCID_50_ titer. Compound cytotoxicity was assessed in parallel in mock-infected plates by MTS cell viability assay at day 3 or 5 p.i.

### Time-of-compound addition assay

A reported method [48] was used, with minor modifications. Confluent HEL299 cells were infected with HCoV-229E and the compounds were added to separate wells at -0.5, 0.5, 2, 4, 6, 8, 10, 12 or 14 h p.i. At 16 h p.i., the cells were washed with ice-cold PBS, then lysed by adding Cells-to-cDNA II lysis buffer and incubating for 10 min at 75 °C. The viral RNA copy number in each sample was determined by one-step RT-qPCR as explained above. The starting number of viral RNA copies (= input RNA) was determined in a cell lysate (from the no compound condition) made at 2 h p.i.

### Selection of compound-resistant HCoV-229E viruses

Instead of performing serial virus passaging, we used a faster resistance selection method (that was earlier developed for Chikungunya virus by Delang et al. [49]). First, HEL299 cells were exposed to serial dilutions of HCoV-229E and compound (EPB-113 or JMLv-061). At day 4 p.i., the optimal condition for resistance selection was defined as the condition receiving the highest MOI and lowest compound concentration achieving complete prevention of CPE. Subsequently, this virus/compound condition was applied to all wells of a 96-well plate with HEL299 cells. At day 4 p.i., the CPE was scored by microscopy and the wells showing virus breakthrough were selected to harvest supernatants and expand the progeny virus. For virus sequencing, we prepared RNA extracts from the expanded virus stocks (QIAamp Viral RNA Kit; Qiagen) and performed high-fidelity RT-PCR (SuperScript IV One-Step RT-PCR System; Invitrogen), yielding five overlapping cDNA fragments that covered the entire viral genome. After Sanger sequencing (Macrogen Europe) with a set of 39 primers, the mutations selected under EPB-113 or JMLv-061 were identified by aligning the sequences of compound-selected and wild-type (WT) forms of HCoV-229E, using CLC Main Workbench (Qiagen). To conduct antiviral experiments, the mutant viruses were plaque-purified in HEL299 cells grown in 6-well plates in the presence of 20 µM compound and overlaid with 0.4% UltraPure Low Melting Point Agarose (Invitrogen).

### Reverse-engineering on SARS-CoV-2

Starting from seven reverse genetics plasmids which were generously donated by P.-Y. Shi (World Reference Center for Emerging Viruses and Arboviruses, Texas) [50], we first replaced their sequences encoding SARS-CoV-2 strain SARS-CoV-2/USA_WA1/2020 by the analogous sequences of Omicron subvariant BF.7. Therefore, viral RNA was extracted (QIAamp viral RNA kit, Qiagen) from a clinical isolate (pango lineage BF.7; hCoV-19/Belgium/rega-43668/2022; EPI_ISL_14623620), followed by reverse transcription (Superscript IV RT First-Strand synthesis system, Invitrogen) and PCR (Platinum SuperFi PCR, Invitrogen). The resulting fragments were used to replace the WA1-sequences in the seven reverse genetics plasmids (NEBuilder HiFi DNA Assembly Cloning Kit, New England Biolabs). For generating mutants, site-directed mutagenesis was performed with overlapping mutagenic primers and Platinum SuperFi II DNA Polymerase (Invitrogen). The procedure for reverse-engineering SARS-CoV-2 was adapted from Xie et al [50]. The plasmids were transformed into One Shot TOP10 Chemically Competent *E. coli* (Invitrogen; for pUC57-based plasmids) or TransforMax EPI300 *E. coli* (LGC Biosearch Technologies; for pCC1-based plasmids) and purified with the Qiagen Plasmid Plus Maxi Kit. All plasmids were validated by Sanger sequencing (Macrogen Europe). Next, SARS-CoV-2 DNA fragments were recovered through restriction enzyme digestion of the seven plasmids and the subsequent ligation step yielded full-length viral DNA. N-gene cDNA was prepared from the plasmid containing the last part of the viral DNA via PCR (Platinum SuperFI II, Invitrogen). Using T7 polymerase (mMESSAGE mMACHINE T7 transcription kit; Invitrogen), the full-length and N-gene viral DNA were *in vitro* transcribed to obtain genomic and N-coding SARS-CoV-2 RNA, followed by purification by phenol:chloroform extraction and isopropanol precipitation. Twenty µg of each was subsequently electroporated into 8 million BHK-21J cells resuspended in PBS, using the Gene Pulser Xcell Electroporation System (Bio-Rad). The electroporated cells were then seeded in a 75-cm² flask with A549^ACE2/TMPRSS2^ cells. The supernatants were harvested after three days, and further expanded in A549^ACE2/TMPRSS2^ cells. The resulting virus stocks were first subjected to nsp15 sequencing, with the primers described by Xie et al. [50], to ensure that the desired mutations were present.

### HCoV-229E replication kinetics and expression of innate immune factors

HEL299, 16HBE, Huh7-STAT1-KO and human macrophage cells were infected with WT or nsp15-mutant HCoV-229E and 2 h later, the inoculum was replaced by fresh medium. At 2, 24, 48 and 72 h p.i., supernatant samples were collected and the cells were extracted for total RNA (ReliaPrep RNA miniprep system, Promega). The supernatants were analyzed for number of viral RNA genomes, using the one-step RT-qPCR method from above, as well as for bioactive IFN-α/β (see below). To measure the mRNA levels for a selection of innate immune factors, the cellular RNA extracts were submitted to RT-qPCR (High-Capacity cDNA Reverse Transcription Kit, Applied Biosystems), using random primers for cDNA synthesis and gene-specific primers (primer sequences are listed in S2 Appendix) for qPCR (GoTaq qPCR kit, Promega). The transcript levels for innate immune factors were normalized to the geometric mean of those for the housekeeping genes ACTB and HPRT (ΔC_T_) and the fold change was calculated as 2^-ΔΔCT^ over mock (ΔC_T_ target - ΔC_T_ mock).

The levels of bioactive IFN-α/β in the supernatants were quantified in HEK-blue IFN-α/β reporter cells, following the manufacturer’s instructions. In brief, 20 µl supernatant was added to the wells of a 96-well plate and the reporter cells were added at 50,000 cells in a volume of 180 µl. Recombinant human IFN-β was included as a positive control. After overnight incubation at 37 °C, 20-µl supernatant samples were collected from the reporter cells, then combined with 180 µl QUANTI-blue solution (InvivoGen) in a 96-well plate and incubated for 2 h at 37 °C. The A_620_ values were measured in a Tecan Spark 10M plate reader. For calculations, all absorbance values were first adjusted by subtracting the blank, i.e. QUANTI-blue in the corresponding medium per cell line, then expressed as the fold increase for virus-versus mock-infected cells.

To assess viral growth at different temperatures, the stocks of WT and mutant viruses were serially (1/10) diluted and incubated in quadruplicate with HEL299 and Huh7-STAT1-KO cells at 33, 35, 37 or 38 °C. After scoring the CPE at day 5 (HEL299) or day 3 (Huh7-STAT1-KO), the CCID_50_ titers were calculated by the method of Reed and Muench.

### Competitive fitness of nsp15 inhibitor-resistant viruses

HEL299 cells were co-infected with WT and nsp15-mutant HCoV-229E virus, mixed at a ratio of 9:1, 1:1 or 1:9. After three days incubation, the supernatants were harvested, titrated and applied to fresh HEL299 cells. This procedure was repeated for an extra three passages. The supernatants of each passage were collected for RNA extraction (QIAamp Viral RNA Kit, Qiagen) followed by RT-PCR (SuperScript IV One-Step RT-PCR, Invitrogen) with primers 229E-For17743 and 229E-Rev24484 (sequences provided in S2 Appendix), resulting in ∼6.7 kb amplicons covering the nsp15 gene. All PCR products were sequenced using Oxford Nanopore Technologies devices. For passages 1 and 2, library preparation was performed using the SQK-RBK110.96 kit and sequencing was done on an r9.4.1 flow cell connected to a GridION (MinKNOW v23.04.6; super-accurate basecalling). For passages 3 and 4, library preparation was done using the SQK-NBD114.96 kit and the resulting library was sequenced on an r10.4.1 PromethION flow cell using a P2 Solo device connected to a GridION (MinKNOW v23.07.12; super-accurate basecalling). Filtlong (v0.2.1) was used to subset for each barcode the 10% highest quality reads with a length between 6.5 and 7 kb. These subsets were mapped against the HCoV-229E reference sequence (NC_002645.1) using the minimap2 (v2.24) map-ont preset [51], resulting in a minimal coverage of 1200X at each site. Detailed coverage statistics per position were determined using bam-readcount (v1.0.1), after which a custom bash script was used to calculate simplified percentages of WT and nsp-15 mutant motif prevalence at each virus passage [52].

### Production of recombinant nsp15 protein from HCoV-229E and SARS-CoV-2

The pET28a(+) plasmid for bacterial expression of SARS-CoV-2 nsp15 was a kind gift from B. Canard (Aix-Marseille University). To engineer the corresponding plasmid for HCoV-229E nsp15, the sequence (codon-optimized for expression in *E. coli*) was supplemented with an N-terminal octahistidine tag and TEV cleavage site and ordered as a gBlock from IDT. This DNA was cloned in the pET28a(+) vector using the NEBuilder HiFi DNA assembly cloning kit (NEB) and corresponding instructions. Starting from this WT-229E nsp15 sequence, the G248V, H250Y and H250A mutants (numbering based on the virus sequence) were created by site-directed mutagenesis with overlapping mutagenic primers and Platinum SuperFi II DNA Polymerase (Invitrogen). To produce the nsp15 proteins, T7 Express Competent *E. coli* cells (NEB) were transformed with pET28a(+)-nsp15 plasmid, as instructed by the supplier. A starter culture was grown overnight in 100 ml Luria–Bertani (LB) broth with 50 µg/ml kanamycin, using a shaking incubator (100 rpm) at 37 °C. From this, 10 ml was added to 1 liter LB medium with 50 µg/ml kanamycin, and the bacteria were grown until an OD_600_ of ∼0.6 was reached. Protein expression was induced at 500 µM isopropyl β-D-1-thiogalactopyranoside, and the cultures were incubated overnight at 18 °C, after which the cells were pelleted by centrifugation (30 min, 5000 g, 4 °C) and stored at -20 °C. For protein purification, the pellets were resuspended in lysis buffer (50 mM Tris pH 8; 300 mM NaCl; 20 mM imidazole; and 1 mM dithiothreitol (DTT), supplemented with 1 mM phenylmethylsulfonyl fluoride, 1% Triton-X, 30 mg lysozyme, 20 mM MgSO4, 100 U DNase I). After cell disruption by high pressure using EmulsiFlex-C3 (Avestin), 4 times at 20,000 bar, cell debris was removed by centrifugation (40 min, 5000 x g, 4 °C) and filtration through a 0.45 µm Whatman GD/XP Syringe Filter (Cytiva). The filtered lysates were subjected to Ni-NTA chromatography (Qiagen) equilibrated with the lysis buffer without supplement, washed and eluted with the increasing concentration of the imidazole (20-300 mM Imidazole). The eluted protein is then applied to a HiLoad Superdex 16/600 200 pg size-exclusion chromatography (SEC) column (GE Healthcare) and run in a buffer containing 25 mM HEPES at pH 8, 300 mM NaCl, 10% glycerol, and 1 mM tris(2-carboxyethyl)phosphine. Finally, the protein solutions were concentrated with a centricon (VivaSpin 15R centricon, 10,000 MWCO; Sartorius AG). The protein products were analyzed by dynamic light scattering (DLS; S2 Fig), which showed that the nsp15 proteins were ≥ 98% pure and present in hexameric form, in line with other reports [53, 54].

### Biophysical evaluation of the compound-nsp15 interaction

The thermal shift assay was executed in white PCR plates. The wells contained 6 µM nsp15 (diluted in assay buffer: 20 mM HEPES pH 7.5; 50 mM NaCl; 5 mM MnCl_2_; and 1 mM DTT), 5x SYPRO orange dye and either 100 µM test compound or the corresponding percentage of DMSO. Using a QuantStudio 5 Real-Time PCR system (Thermo Fisher), fluorescence was continuously monitored while the plate was heated from 4 to 95 °C at a ramp rate of 0.05 °C/s. The nsp15 melting temperature (T_m_) was calculated by determining the temperature at which the negative first derivative of the fluorescence versus temperature (-dFluorescence/dT) reaches its minimum, using Protein Thermal Shift Software v1.3 (Thermo Fisher).

To assess, by surface plasmon resonance (SPR), the interaction between EPB-113 and nsp15 from HCoV-229E (WT or mutant) or SARS-CoV-2 we used a Biacore 8K instrument (Cytiva). Nsp15 was covalently coupled onto a CM4 sensor chip (Cytiva) at a density of around 3500 resonance units (RU) using standard amine coupling chemistry in 10 mM acetate buffer pH 4.5. A reference flow was used as a control for non-specific binding and refractive index changes. All binding experiments were performed at 25 °C in 50 mM Tris buffer supplemented with 500 mM NaCl, 5 mM MnCl_2_, 0.05% Tween20 and 5% DMSO. Serial dilutions of EPB-113 or EPB-102 were injected for 120 s at a flow rate of 30 μl/min. The dissociation was monitored for 6 min. Buffer blanks were used for double referencing and a DMSO solvent correction was included. Biacore Insight Software 5.0.18 was used for analysis.

### Enzymatic EndoU assay

For the FRET-based EndoU cleavage assay, we employed a 5-nt single-stranded DNA-RNA hybrid substrate (5′-6-FAM-AArUAA-3′-IABkFQ; synthesized by IDT), that contained a single U®A-cleavage site, 5′-carboxyfluorescein (FAM) fluorophore and 3′-Iowa Black quencher [29]. The assay was conducted in black half-area 96-well plates, using the same assay buffer as in the thermal shift assay. The wells (25 µl total volume) contained 63 nM (HCoV-229E) or 4 nM (SARS-CoV-2) nsp15, 0.5 µM substrate and 100 µM of compound or the corresponding percentage of DMSO. Using a Tecan Spark 10M plate reader set at 485 nm emission and 535 nm excitation wavelength, the increasing fluorescence was continuously measured during 1 h at 37 °C. The initial cleavage rate (*V_0_*) was obtained by calculating the slope (RFU/s) of the 5-10 min time interval of the cleavage reaction.

In parallel, the samples after 1 h reaction time were analyzed by HPLC on a capillary chromatography instrument (CapLC, Waters) equipped with a C18 reversed-phase column (PepMap 0.5 x 15 mm, LC Packings). The elution buffer contained 0.05% v/v *N,N*-dimethylaminobutane (DMAB, Acros) as ion pairing reagent and 1% v/v 1,1,1,3,3,3-hexafluoro-2-propanol (hexafluoroisopropanol, HFiP, Acros) in water (pH 8.0). The organic phase consisted of 50% v/v acetonitrile (Fisher Scientific) in water. Oligonucleotides were eluted at a flow rate of 12 μl per min, applying a gradient starting at 5% organic phase and increasing by 20% per min during 4 min. The mass spectrometer consisted of a quadrupole time-of-flight instrument (Synapt G2 HDMS, Waters) equipped with a standard electrospray source. Every 2 s, spectra were recorded in negative ion mode with following settings: cone voltage 35 V, capillary voltage 2200 V, source temperature 80 °C and N_2_ desolvation gas 600 L/h. The acquired spectra were mass corrected using leucine enkephalin and deconvoluted applying the MaxEnt algorithm (MassLynx 4.1, Waters).

### Data analysis and statistics

All graphs were created with GraphPad Prism 10.3.1 software, which was also used for statistical analysis. Statistical significance is shown as: \*\*\*\**P* ≤ 0.0001, \*\*\**P* ≤ 0.001, \*\**P* ≤ 0.01, \**P* ≤ 0.05, ns: not significant.

## Results

### EPB-113 is a strong replication inhibitor of HCoV-229E that also suppresses SARS-CoV-2

HCoV-229E differs from other human CoVs in causing manifest cytopathic effect (CPE) in different cell lines, while its manipulation is allowed at biosafety level 2. Based on this, we have been using since several years a CPE reduction assay in HCoV-229E-infected HEL299 cells, to conduct convenient phenotypic screening for novel CoV inhibitors. To assess the compounds’ antiviral effect, the viral CPE is quantified by MTS cell viability assay in combination with microscopic examination. These two methods are also used to determine compound cytotoxicity in mock-infected cells.

Our screening campaign identified compound EPB-113 as a robust lead compound, exhibiting an EC_50_ value for HCoV-229E of 0.48 µM and CC_50_ value for HEL299 cells of 92 µM, thus yielding a favorable selectivity index (SI; ratio of CC_50_ to EC_50_) of 193 (Figs 1B and 1D). Antiviral activity was very similar whether determined by MTS assay or by microscopic scoring (Fig 1D). The activity was confirmed when we quantified the virus in the supernatant at day 3 p.i. Here, EPB-113 showed an EC_90_ value of 0.65 µM based on RT-qPCR for viral genome copies and of 0.40 µM based on titration of infectious virus (Figs 1B and 1D). In HCoV-229E-infected HEL299 cells, EPB-113 outperformed GS-441524, the nucleoside form of remdesivir that was included as reference compound [EC_50_ value of 3.5 µM (CPE assay) and EC_90_ value of 2.5 and 4.6 µM (by infectious virus titration and RT-qPCR, respectively)].

**Fig 1.**
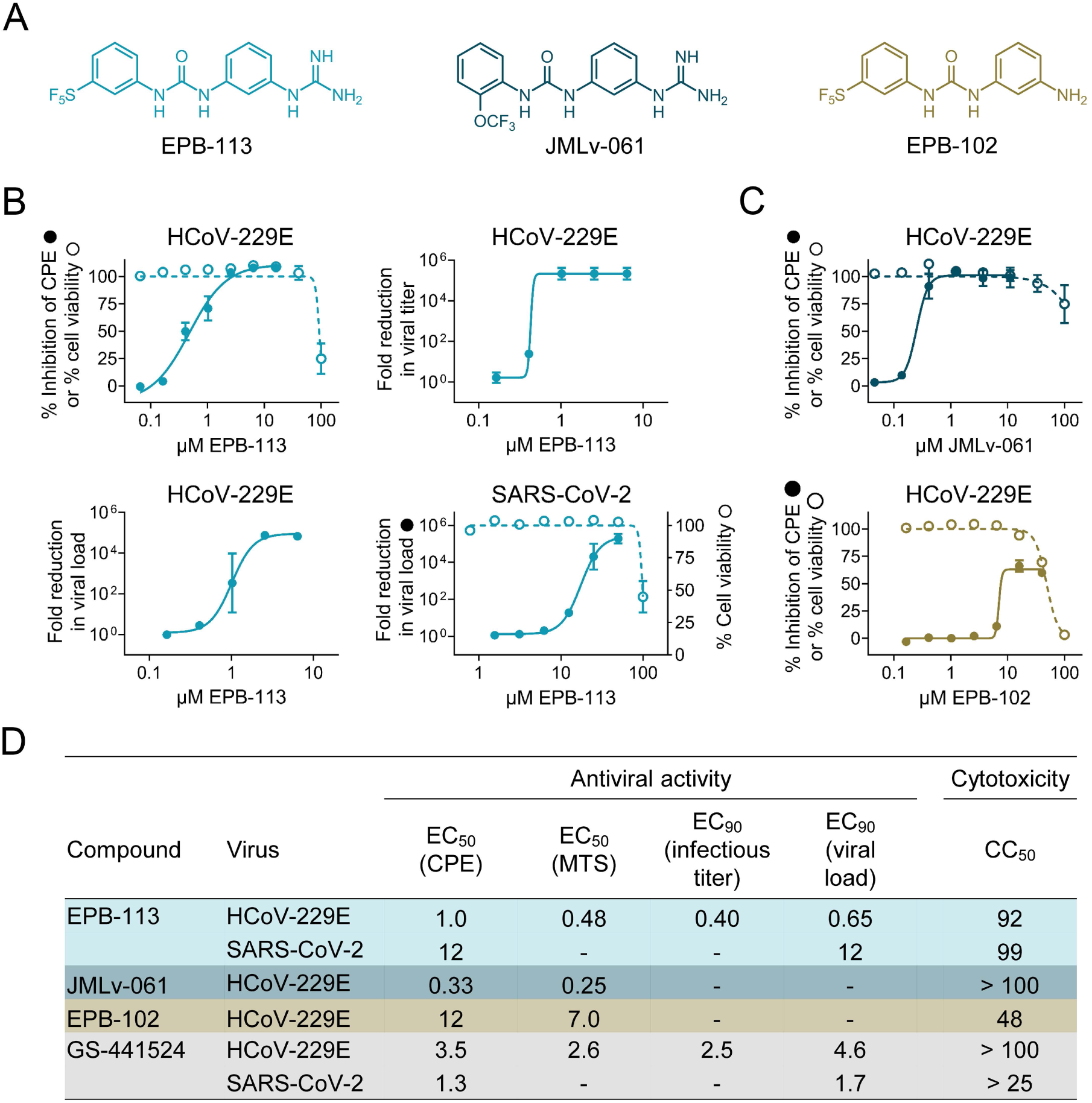
Inhibitory activity of EPB-113 on the replication of HCoV-229E and SARS-CoV-2. (A) Chemical structures of EPB-113 and its structural analogues EPB-102 and JMLv-061. (B) Inhibition of HCoV-229E replication in HEL299 cells was determined by MTS-based CPE reduction assay; infectious virus titration on the supernatant; and RT-qPCR-based viral load assay. Inhibition of SARS-CoV-2 replication in A549^ACE2/TMPRSS2^ cells was quantified by viral load assay. Cytotoxicity was determined by MTS cell viability assay in mock-infected cells and is shown in dashed lines (open circles). Data points are the mean ± SEM (N=3-12). (C) Anti-HCoV-229E activity and cytotoxicity in HEL299 cells for EPB-102 and JMLv-061, two structural analogues of EPB-113. Data points are the mean ± SEM (N=3). (D) Table summarizing the values for antiviral activity and cytotoxicity of EPB-113, JMLv-061, EPB-102 and reference compound GS-441524.

Chemically, EPB-113 is a diphenylurea compound featuring a *m*-guanidine function in an aromatic ring and a *m*-pentafluorosulfanyl group in the other aromatic ring (Fig 1A). To determine the structure-activity relationship for HCoV-229E, we synthesized a series of more than 20 structural analogues, the outcome of which will be published in a separate report (S. Vazquez, manuscript in preparation). Of note, as the very lipophilic *m*-pentafluorosulfanyl group is uncommon in medicinal chemistry, we also evaluated other analogs lacking this group, such as JMLv-061 (Fig 1A), with an *o*-trifluoromethoxy group instead of the *m*-pentafluorosulfanyl unit, but keeping the *m*-guanidine group on the other aromatic ring. While this change had no impact on the antiviral activity (EC_50_ for JMLv-061: 0.33 µM in HCoV-229E infected HEL299 cells; Figs 1C and 1D), the analog lacking the guanidine function (= EPB-102; Fig 1A) was shown to have much weaker activity (EC_50_ of 12 µM in HCoV-229E infected HEL299 cells, reaching only 60% inhibition at 40 µM; Figs 1C and 1D), thus highlighting the relevance of the guanidine group in the antiviral activity.

EPB-113 proved also a robust inhibitor of SARS-CoV-2 replication in A549^ACE2/TMPRSS2^ cells. The molecule was shown to have an EC_90_ value (by RT-qPCR for viral load) of 12 µM and to produce more than 5-log_10_ reduction in viral load at 50 µM of compound (Fig 1B). The cell viability of mock-infected A549^ACE2/TMPRSS2^ cells was 100% at 50 µM EPB-113 and 45% at 100 µM compound (Fig 1B), which is similar to what was seen in HEL299 cells.

### Viral resistance to EPB-113 maps to the catalytic core of nsp15 EndoU

To identify the target of EPB-113, we performed selection of resistant virus. For this and all subsequent mechanistic experiments, we chose to use HCoV-229E instead of SARS-CoV-2, since this allowed to apply the compound at a concentration that was well above the EC_50_ value. In the first step (Fig 2A), we determined the optimal condition, being the highest MOI of HCoV-229E and lowest compound concentration for which viral CPE was completely inhibited. Next, we applied this condition (i.e. 20 µM of compound) to all wells of a 96-well plate with HEL299 cells. To maximize the chances of obtaining resistance, we added EPB-113 to one plate and its analogue JMLv-061 to a second plate. At day 4 p.i., the few wells showing virus breakthrough were selected for virus expansion and subsequent full-genome sequencing.

**Fig 2.**
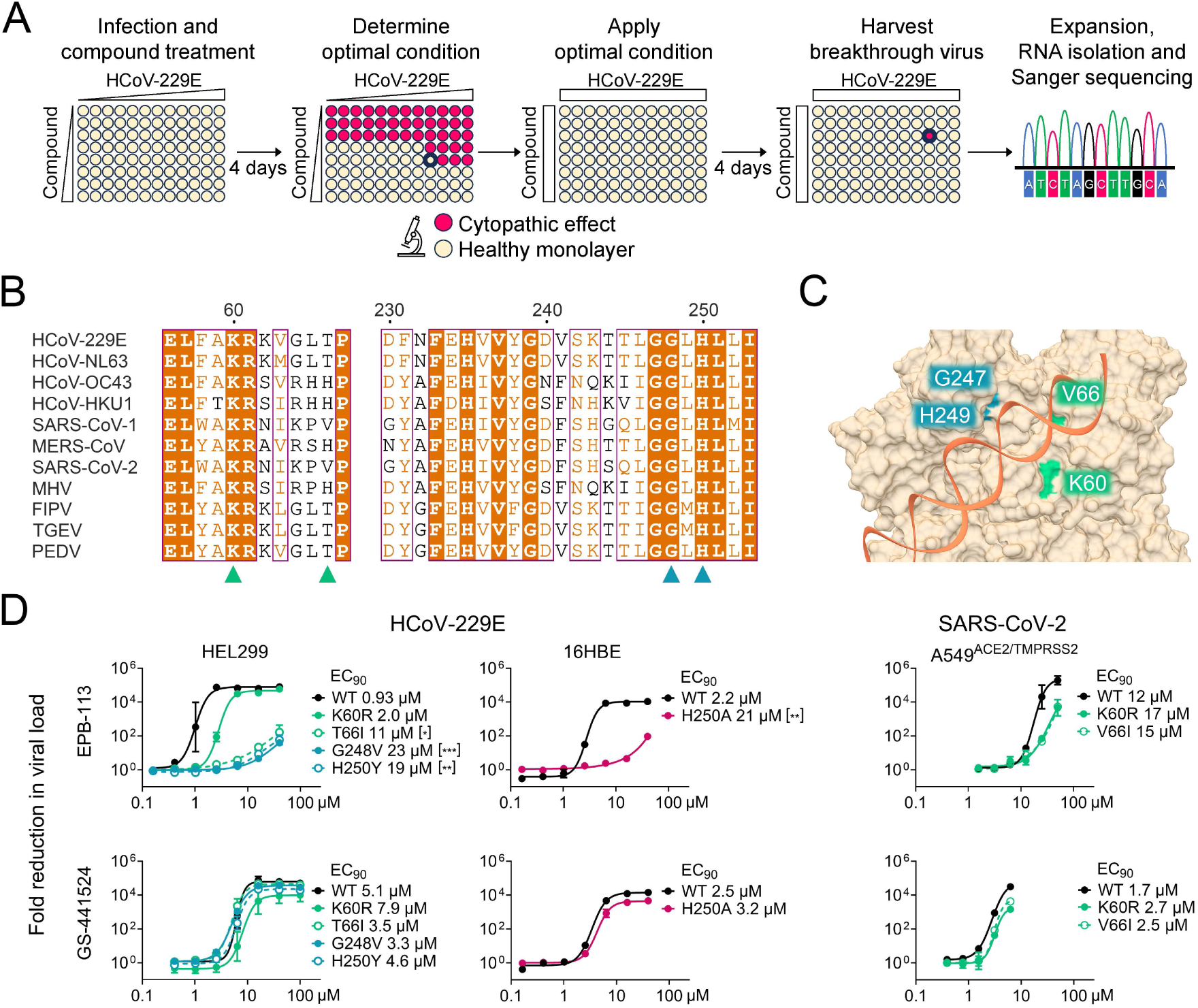
Resistance to EPB-113 maps to the catalytic core of nsp15 EndoU. (A) Setup of the resistance selection experiment. (B) Alignment of partial nsp15 sequences from 11 coronaviruses, with EPB-113 and 5h resistance sites marked with triangles. Both EPB-113 resistance sites, G248 and H250, are located at the catalytic core of nsp15 EndoU and strictly conserved in all CoVs, with H250 being part of the catalytic triad. Sequence similarities from Clustal Omega-aligned sequences were rendered using ESPript 3.0. (C) Structure of the catalytic core of SARS-CoV-2 EndoU bound to dsRNA (PDB 7TJ2) [42]. Resistance of HCoV-229E to EPB-113 maps to residues G248 and H250, while that for 5h maps to K60 and T66 (corresponding to residues K60, V66, G247 and H249 in SARS-CoV-2). (D) Evaluation of EPB-113 and reference compound GS-441524 against nsp15-mutant CoVs, based on RT-qPCR-based viral load assay. Left panels: HEL299 cells infected with HCoV-229E WT; the G248V_nsp15_ or H250Y_nsp15_ mutant (obtained in this study); or the K60R_nsp15_ or T66_nsp15_ mutant (previously selected under compound 5h) [42]. Middle panels: 16HBE cells infected with H250A_nsp15_ mutant HCoV-229E, created by reverse genetics [11]. Right panels: A549^ACE2/TMPRSS2^ cells infected with K60R_nsp15_ or V66I_nsp15_ mutant SARS-CoV-2, created by reverse genetics. Data points are the mean ± SEM (N=2-3). The panel legends show the EC_90_ values and statistical significance of the difference between mutant and WT, based on (left panels) ordinary one-way ANOVA with Dunnett’s corrections for multiple comparisons and (middle panels) two-tailed unpaired t-test.

The breakthrough viruses were found to carry mutations H250Y and G248V in the EndoU domain of nsp15, after selection under EPB-113 and JMLv-061, respectively. After obtaining plaque-purified mutant viruses, we confirmed their resistance by RT-qPCR-based viral load assay, in which EPB-113 showed an increase in EC_90_ value between mutant and WT of 20- to 25-fold (Fig 2D; we no longer included JMLv-061 since its antiviral activity for WT was very similar to that of EPB-113). Also another variant at the His250 site, i.e. the H250A_nsp15_ mutant form of HCoV-229E which was created by the V. Thiel team using reverse-engineering [11], proved to be highly resistant to EPB-113 (Fig 2D). In this case the assay was done in 16HBE cells, since the H250A_nsp15_ virus showed significantly impaired replication in HEL299 cells (see below). The reference compound GS-441524 was equally active against the nsp15-mutant viruses and WT.

Residues Gly248 and His250 are strictly conserved among CoVs and located in the catalytic center of EndoU (Figs 2B and 2C). While His250 is part of the catalytic triad, the nearby Gly248 residue was reported to engage in hydrogen bonding with the 3′-phosphate of the cleavage product [36].

As mentioned in the introduction, our group previously [42] identified betulonic acid derivative 5h as a strong inhibitor of HCoV-229E which selects for resistance mutations K60R_nsp15_ and T66I_nsp15_ near the EndoU catalytic core (Fig 2C). Considering that EPB-113 and 5h have totally different chemical structures, we investigated whether the two compounds are synergistic when combined. Data analysis with the zero-interaction potency (ZIP) model in SynergyFinder+ (S3 Fig) [55] indicated that the combination of EPB-113 and 5h is synergistic (mean ZIP synergy score: 12), suggesting that these two inhibitors possess non-overlapping mechanisms that potentiate each other. The combinations of EPB-113 plus GS-441524 or 5h plus GS-441524 proved additive (mean ZIP synergy score below 10 in both cases).

We next investigated potential cross-resistance between the two inhibitor classes. For this experiment we used mutant viruses T66I_nsp15_ and K60R_nsp15_, which arose after virus passaging under 5h [42]. Fig 2D shows that the activity of EPB-113 was strongly affected by T66I_nsp15_, the shift in EC_90_ value versus WT virus being almost equally pronounced (12-fold) as for G248V_nsp15_ and H250Y_nsp15_. There was also a 2-fold, yet non-significant, shift for the K60R_nsp15_ mutant.

To assess whether the above resistance profile of HCoV-229E also applies to SARS-CoV-2, reverse engineering was performed to introduce the above mutations in SARS-CoV-2. We were unable to rescue viable SARS-CoV-2 virus carrying substitution G247V_nsp15_, H249Y_nsp15_ or H249A_nsp15_. Although this observation underscores the importance of a functional EndoU during SARS-CoV-2 replication, we did not expect it, as all three mutations proved clearly viable in the context of HCoV-229E. On the other hand, we successfully generated the K60R_nsp15_ and V66I_nsp15_ mutants of SARS-CoV-2 (corresponding to K60R_nsp15_ and T66I_nsp15_ in HCoV-229E), and both viruses proved to be less sensitive to EPB-113 than WT virus (Fig 2D, right panels).

In short, resistance selection with HCoV-229E, combined with reverse engineering on SARS-CoV-2, showed that the antiviral effect of EPB-113 is tightly linked with nsp15, and specifically its EndoU domain.

### EPB-113 mainly acts in an IFN-independent manner

So far, the main validated function of nsp15 EndoU is to avoid early induction of IFN-I by reducing the levels of viral dsRNA replication intermediates [10–12]. Considering that EPB-113 selects mutations in the EndoU catalytic core, we questioned whether its antiviral activity might depend on the cell’s ability to mount an IFN-I response towards CoV infection. To address this, we determined the anti-HCoV-229E-activity of EPB-113 in human blood-derived macrophages, HEL299 human fibroblast cells (in which our mutants arose under selective pressure of EPB-113 or 5h), 16HBE human bronchial epithelial cells and Huh7-STAT1-KO cells; the latter cannot mount the signaling response downstream to IFN-I binding. As shown in Fig 3, the compound proved nicely active in all four cell types, the EC_90_ values being in the range of 0.14-2.5 µM. The strong activity in the IFN-I non-responsive Huh7-STAT1-KO cells attracted our special attention since it indicated that EPB-113 does not have a major dependence on IFN-I to suppress viral replication.

**Fig 3.**
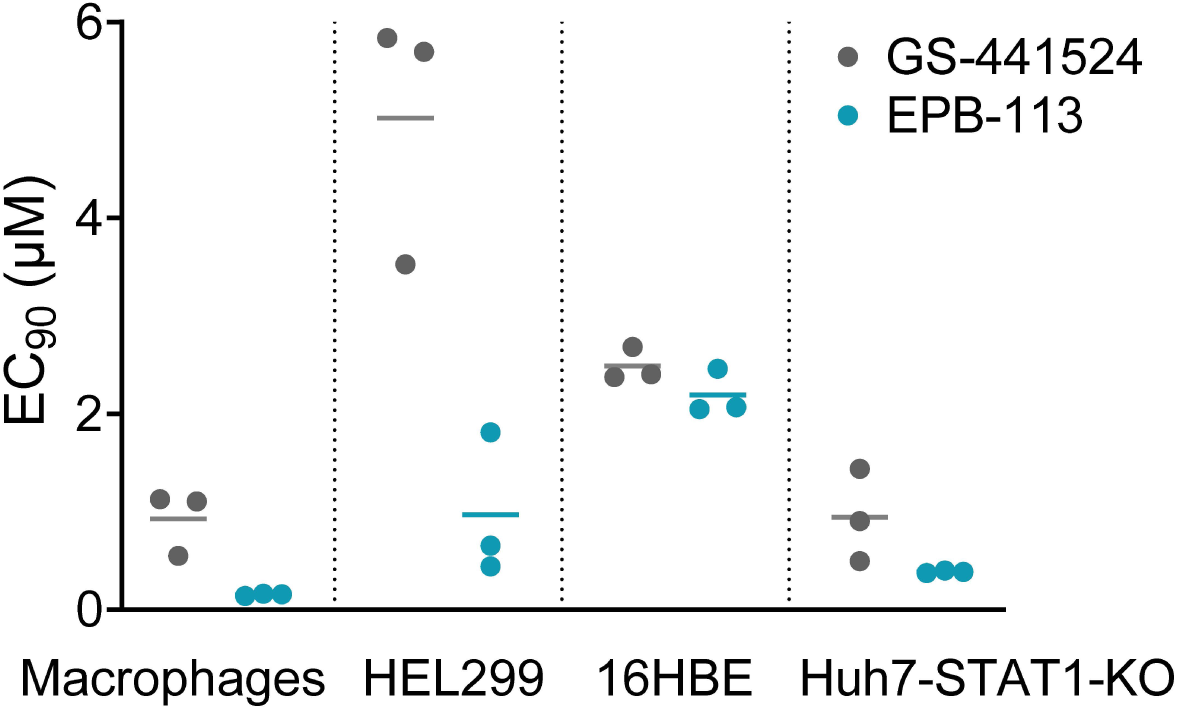
EPB-113 is active in both IFN responsive and non-responsive cell types. EC_90_ values of EPB-113 and GS-441524 for inhibition of HCoV-229E WT virus in human macrophages, HEL299 cells, 16HBE cells and Huh7-STAT1-KO cells, based on viral load assay. The graph shows the individual values and mean from three independent experiments.

### EPB-113 acts at a post-entry and intermediate stage in first-cycle HCoV-229E replication

Since the above findings provided strong support for a direct (i.e. IFN-independent) antiviral effect of EPB-113, we wanted to gain basic insight into which stage of the replication cycle could be affected. To address this, we conducted one-cycle time-of-addition experiments in HCoV-229E-infected HEL299 cells. Besides EPB-113, we included four reference compounds: E64d (an inhibitor of cathepsin-mediated entry of the HCoV-229E ATCC strain) [56]; nirmatrelvir (an M^pro^ inhibitor); GS-441524 (an RdRp inhibitor that terminates viral RNA synthesis); and K22 (which prevents nsp6-mediated formation of the double-membrane vesicle structures where viral RNA synthesis takes place) [57]. The compounds were added to the cells at either 30 min before or several time points after virus infection. At 16 h p.i. (previously determined to mark the end of one HCoV-229E replication cycle), the cells were collected to quantify the level of intracellular viral RNA by RT-qPCR. We also analyzed a condition at 2 h p.i. (= RNA from input virus) to express the data as the fold increase in viral RNA at 16 h p.i. versus input. In the absence of compound, this fold increase was ∼10,000-fold (= factor of 4 log_10_ in Fig 4).

**Fig 4.**
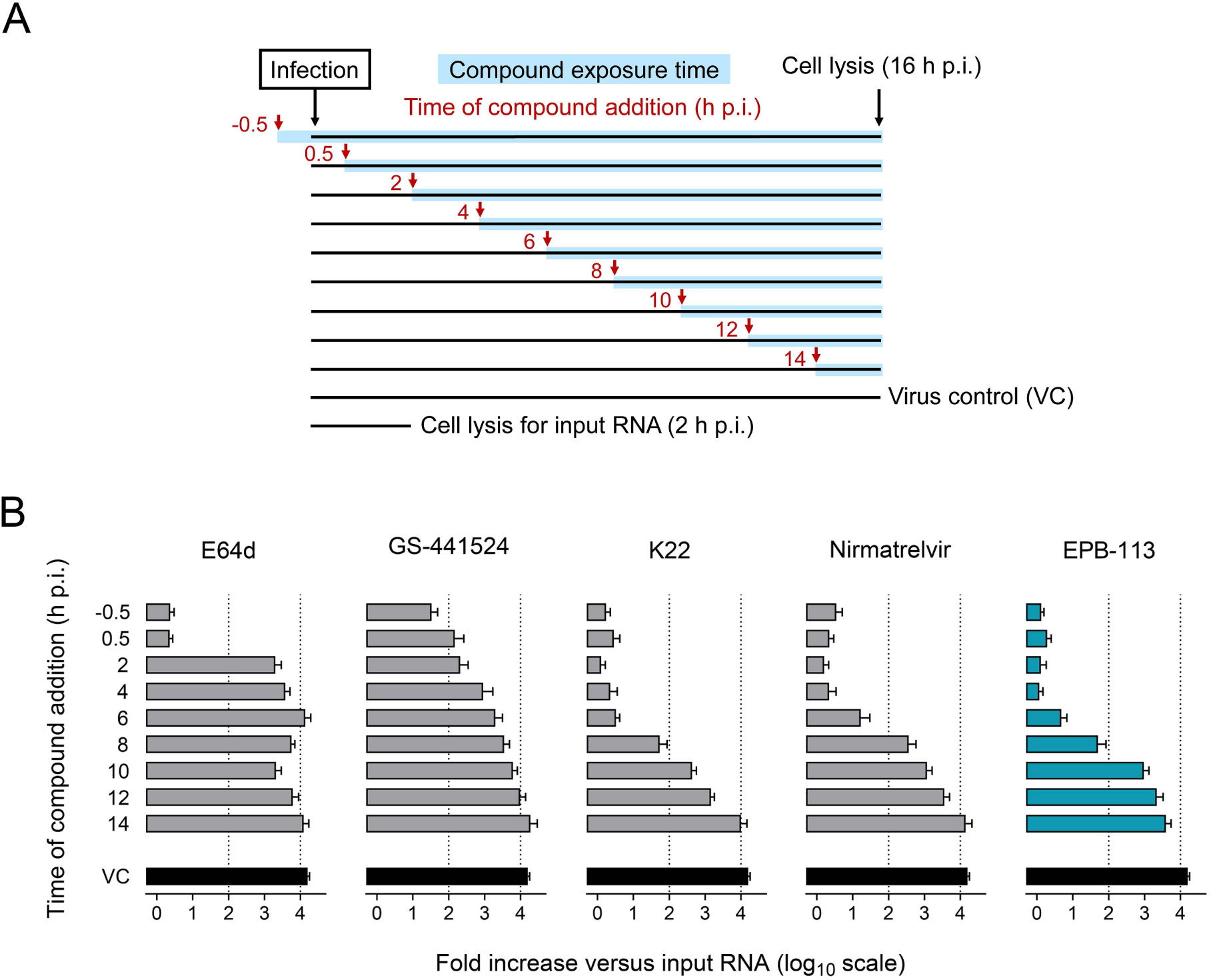
Action point of EPB-113 and reference compounds in a one-cycle time-of-addition assay. (A) Experimental setup. (B) Timewise, EPB-113 has a similar action point as K22, located at an intermediate stage in HCoV-229E replication. The fold increase in intracellular viral RNA is expressed relative to the amount of input RNA at 2 h p.i. Compound concentrations: 15 µM E64d; 15 µM GS-441524; 15 µM K22 [57]; 3 µM nirmatrelvir; and 15 µM EPB-113. Data points are the mean ± SEM (N=3).

Whereas the entry inhibitor E64d lost all of its activity when added at 2 or more hours p.i., EPB-113 and K22 still had profound effect on the level of viral RNA (= ∼300 times less than in the virus control) when added at 8 h p.i. (Fig 4). Nirmatrelvir already started to lose activity at 8 h p.i. (only 40-fold reduction versus virus control), consistent with inhibition of protease activity. GS-441524 was already less active when added at 4 h p.i., indicating that this nucleoside analogue requires a few hours to attain a sufficient level of its active 5′-triphosphate metabolite in HEL299 cells. The experiment further showed that neither of the tested compounds, including EPB-113, act at the stages of virus maturation or budding. Inhibitors of the latter processes would show no inhibition in our assay, since we measured the viral RNA levels inside the cells and not the progeny virus released in the supernatant. Besides, the data corroborate that EPB-113 also interferes with viral replication in a direct manner that does not involve IFN-I. The IFN-β mRNA only started to increase at 48 h p.i. of HEL299 cells (see below). Since the level of viral RNA, produced within one replication cycle of ∼16 h, is too low to induce IFN-I, the strong activity of EPB-113 in the time-of-addition assay must be independent of this cytokine.

### EPB-113 generates a subtle boost in IFN-I, but only when its addition is postponed

To answer whether IFN-I still contributes to the antiviral activity, we assessed whether EPB-113 changes the expression of IFN-β in HCoV-229E-infected HEL299 cells. When the compound was added together with virus (= at 0 h p.i.), the levels of IFN-β mRNA (assessed at 48 h p.i.) were similar in cells receiving no compound, EPB-113, GS-441524 or nirmatrelvir (S4 Fig, panel A). Strikingly, when compound addition was postponed until 24 h p.i. (Fig 5A), which gave the virus the chance to start replicating, we observed 48 h later a significant spike in IFN-β in the condition receiving 1 µM of EPB-113 (Fig 5C). The boost in IFN-I required the presence of virus, since it was not seen in uninfected cells exposed to EPB-113 for 48 h (S4 Fig, panel B). The spike was evident from both IFN-β mRNA quantification and IFN-I bioassay. For this delayed treatment schedule, the dose-response curve showed an extra plateau at exactly this concentration of EPB-113 (Fig 5B). The biphasic shape of the curve indicates a dual antiviral mechanism, in which the compound’s direct antiviral effect combines with an indirect effect through boosting of IFN-I. While GS-441524 had no such effect when added at 24 h p.i., the M^pro^ inhibitor nirmatrelvir effectuated a strong and dose-dependent increase in the levels of IFN-β mRNA and IFN-I protein. The IFN-I increase in the presence of nirmatrelvir implies that, besides its role in polyprotein cleavage, the M^pro^ of HCoV-229E has immune-evading activity, as already demonstrated for the M^pro^ enzymes of MERS-CoV [58] and SARS-CoV-2 [59]. M^pro^ was shown to exhibit cleavage activity towards different cellular proteins [60], including RIG-I [59] and other components of the innate immune response [61]. For both EPB-113 and nirmatrelvir, delaying treatment until 24 h p.i. was essential to ensure the accumulation of sufficient viral RNA to function as a pathogen-associated molecular pattern (PAMP). Subsequently, inhibiting the immune evasion activities of either nsp15 or M^pro^ led to a significant increase in interferon levels in response to the pre-existing viral RNA. Since it has been reported that EndoU-deficient virus triggers apoptotic cell death [10], we evaluated whether EPB-113 might influence this process, which we assessed from cleavage of PARP1, a substrate of caspase-3. Rather than enhancing PARP1 cleavage, EPB-113 inhibited apoptosis under our assay conditions (S5 Fig), which is plausibly explained by its inhibitory effect towards viral CPE.

**Fig 5.**
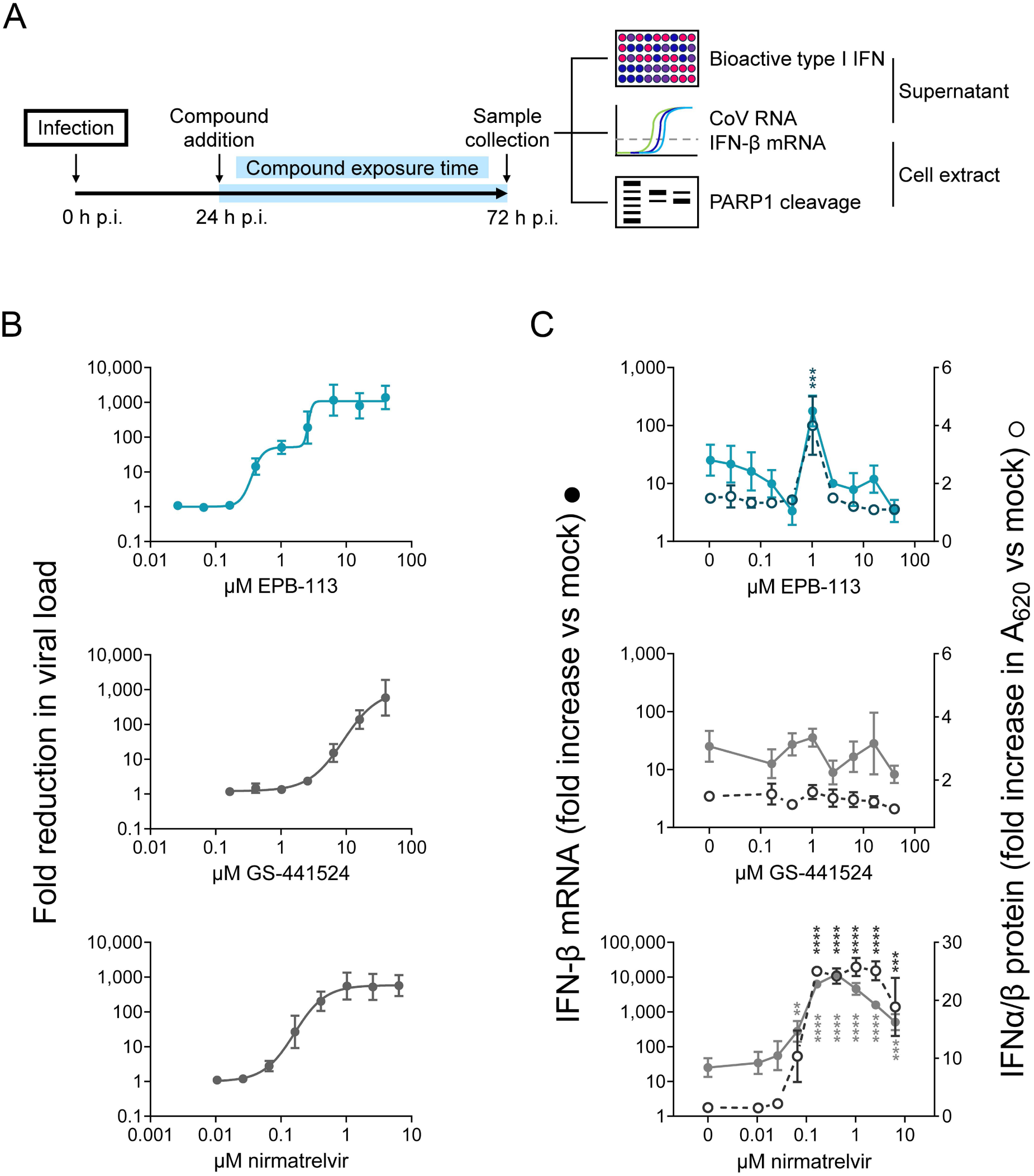
Delaying EPB-113 exposure results in subtle induction of IFN-I. (A) Compound addition to HEL299 cells infected with HCoV-229E WT was delayed until 24 h p.i. and 48 h later, samples were collected for assessment of (B) viral load and (C) levels of IFN-I (left Y-axis: IFN-β mRNA; right Y-axis: bioactive IFN-I). GS-441524 and nirmatrelvir were included as reference compounds. Data points are the mean ± SEM (N=4). Statistical analysis was performed using a one-way ANOVA with Dunnett’s correction for multiple comparisons, to compare each compound concentration to untreated virus control (0 µM).

Taken together, the above results demonstrate that EPB-113 mainly uses an IFN-independent mechanism to suppress HCoV-229E replication at the stage of viral RNA synthesis. This is evident from the prominent activity in Huh7-STAT1-KO cells and strong suppression of first-cycle viral RNA synthesis. In addition, EPB-113 effectuates subtle boosting of IFN-I when treatment is postponed to a later time point. Under these circumstances, the compound has a dual effect, as visible from the biphasic dose-response curve.

### The binding of EPB-113 to nsp15 destabilizes the protein and stimulates its EndoU activity

In the next part, we used different biophysical and biochemical methods to study the binding interaction between nsp15 protein and EPB-113, and how this binding may change the EndoU activity of nsp15. The HCoV-229E nsp15 protein (either WT or the G248V-, H250Y- or H250A-mutant) and SARS-CoV-2 analogue were expressed in *E. coli* as described by others [7, 35] and the hexameric proteins were purified by size-exclusion chromatography. Dynamic light scattering (DLS) analysis confirmed that all five proteins were ≥98% hexameric (S2 Fig).

Using a thermal shift assay, we found that EPB-113 causes quite pronounced reduction in the thermostability of nsp15, producing a shift in the melting temperature (T_m_) of 4°C (Fig 6A). The shift was comparable for the WT proteins of HCoV-229E and SARS-CoV-2. It required a fairly high concentration of 100 µM EPB-113, which represents a 17-fold molar excess compared to the concentration of 6 µM for nsp15 protein. The thermal shift was not at all observed with amine EPB-102, the analogue lacking the guanidine function that exhibited much weaker anti-HCoV-229E activity in HEL299 cells (15-fold higher EC_50_, see first part of the results). The binding of EPB-113 to nsp15 was confirmed by surface plasmon resonance, in which we immobilized nsp15 to the chip as ligand, then used EPB-113 and EPB-102 as analytes. The sensorgrams (Fig 6B) show dose-dependent binding of EPB-113 to nsp15, with a similar profile for the HCoV-229E and SARS-CoV-2 proteins. The binding was markedly less with EPB-102 (5- to 8-fold less RU at 100 µM). Importantly, the nsp15 binding mode of EPB-113 involves the catalytic core of the EndoU domain, since H250A- and H250Y-mutant nsp15 proteins showed little to no binding of the compound, and also mutation G248V had dramatic impact (Fig 6C).

**Fig 6.**
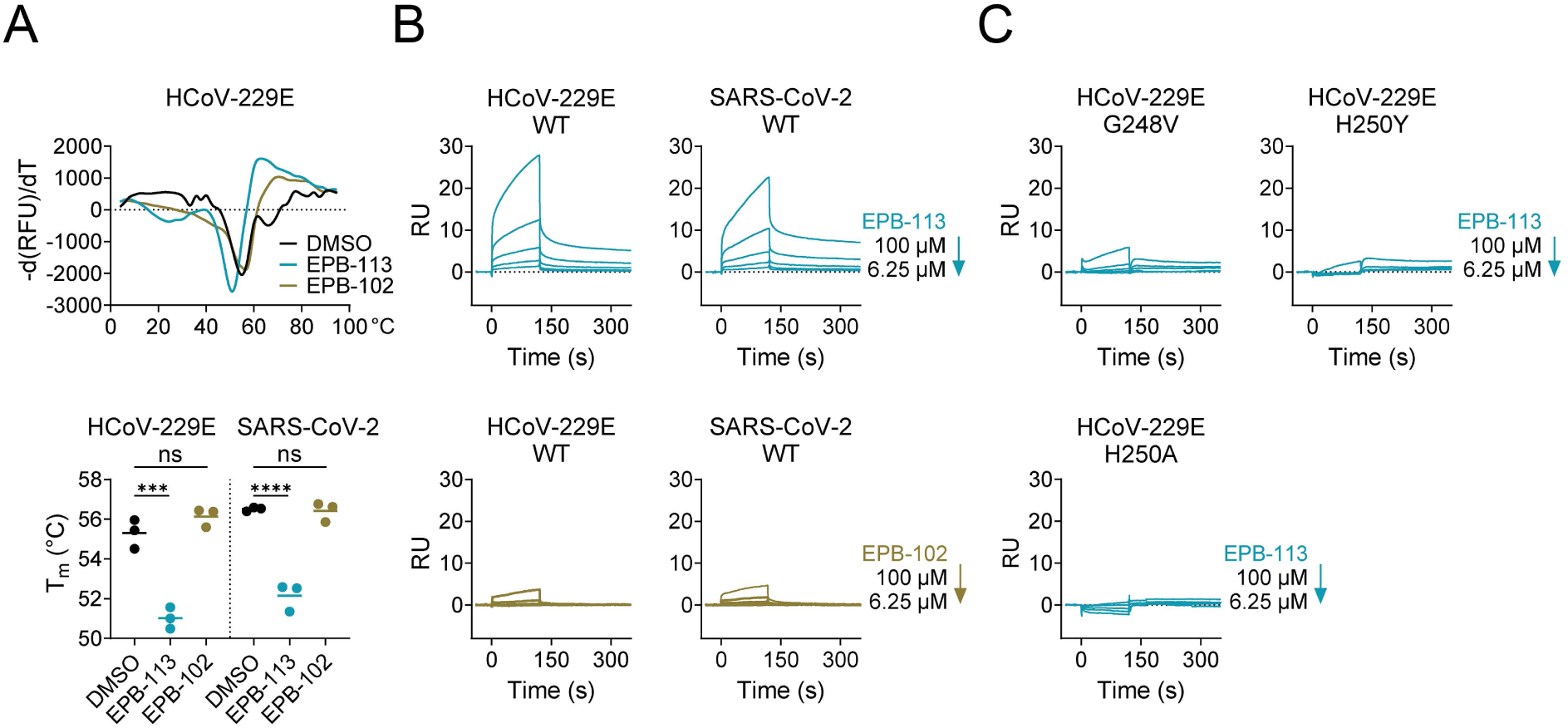
EPB-113 binds to hexameric nsp15 protein. (A) Thermal shift assay showing the decrease in T_m_ when 6 µM of nsp15 protein was incubated with 100 µM EPB-113. The controls contained 100 µM EPB-102 or the equivalent amount (= 5%) of DMSO. The top panel shows the curves generated from the mean data of three independent experiments. The graph (bottom panel) shows the melting temperatures (individual and mean values) in three independent experiments. Statistical significance was determined with an ordinary one-way ANOVA with Dunnett’s correction. (B,C) Assessment of binding by surface-plasmon resonance. RU: resonance units. Panel B: dose-dependent binding between EPB-113 (top panels) or EPB-102 (bottom panels) and recombinant nsp15 from HCoV-229E (left panels) or SARS-CoV-2 (right panels). Panel C: binding of EPB-113 to HCoV-229E nsp15 was severely affected by the G248V-, H250Y- and H250A mutations.

To next investigate the impact of EPB-113 on EndoU cleavage, we conducted a convenient fluorescence resonance energy transfer (FRET) assay that allows to follow the reaction (Fig 7A) in real-time. Quite intriguingly, EPB-113 was found to enhance the EndoU cleavage reaction (Fig 7B). At 100 µM EPB-113, the initial reaction velocity (V_0_) was increased by about 2-(SARS-CoV-2) or 4-fold (HCoV-229E) compared to the DMSO control condition (P ≤ 0.001). Also this effect was not observed with EPB-102. The activating effect of EPB-113 was also evident when we conducted HPLC-MS analysis on the mixtures after 1 h reaction time. This method enabled to discriminate between the starting substrate; 5′-hydroxylated reaction product; 2′,3′-cyclic phosphate reaction intermediate; and 3′-phosphorylated reaction product (see reaction scheme in Fig 7A). The latter two cannot be distinguished in the FRET assay, since they both carry the fluorophore. Although care should be taken when interpreting HPLC-MS peak areas in a quantitative manner, the chromatograms (Fig 7C) show that, in the DMSO control and EPB-102 samples, the 5′-hydroxylated product and 2′,3′-cyclic phosphate intermediate attained only low levels after 1 h reaction time. The minute amount of the 3′-phosphorylated product confirms a previous report that the 2′,3′-cyclic intermediate undergoes slow hydrolysis to this 3′-end product [35]. In the EPB-113 sample, both 5′- and 3′-end products as well as the 2′,3′-cyclic intermediate showed markedly higher peak areas, confirming that the compound stimulates the reaction. Together, these data corroborate that EPB-113 induces prominent activation of the cleavage reaction, while not affecting the relative abundance of the cyclic intermediate and final 3′-phosphate product.

**Fig 7.**
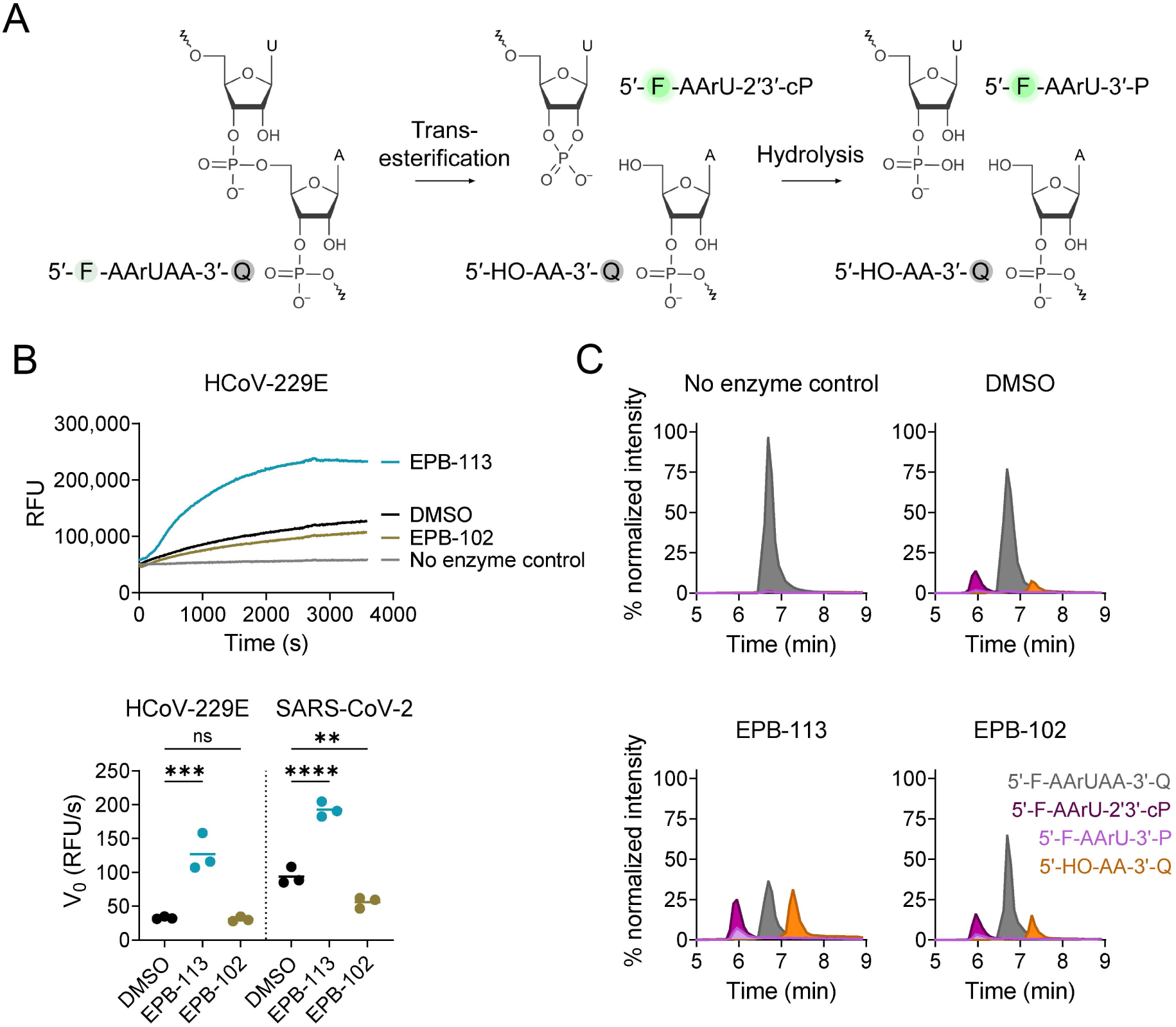
EPB-113 enhances the EndoU cleavage reaction. (A) Scheme of the two-step EndoU cleavage reaction. (B) FRET assay in the presence of 100 µM EPB-113, 100 µM EPB-102 or the equivalent amount (= 5%) of DMSO. RFU: relative fluorescence units. An example of the real-time fluorescence curves is shown in the top panel, while the graph in the bottom panel shows the initial reaction velocity (V_0_) derived from the 5-10 min interval. Individual and mean data from N=3. (C) HPLC-MS analysis on samples collected after 1 h reaction time. In the chromatograms, the colors indicate the FRET substrate; 2′,3′-cyclic intermediate; 5′-hydroxylated product; and 3′-phosphorylated product. The Y-axis shows the % normalized intensity.

### EPB-113-resistant HCoV-229E mutants have a replication deficit

#### Attenuation in human macrophages

In the final part, we addressed whether the replication fitness of HCoV-229E is affected by the nsp15 mutations which cause reduced sensitivity to EPB-113 and are located at (i.e. G248V_nsp15_ and H250Y_nsp15_) or close to (i.e. K60R_nsp15_ and T66I_nsp15_) the EndoU catalytic core. To thoroughly profile these mutants, we compared them to WT virus as well as to the reverse-engineered H250A_nsp15_ virus. The latter was previously reported to be severely attenuated in human macrophages, due to early induction of IFN-I [11]. Hence, we compared viral growth in human macrophage, HEL299, 16HBE and Huh7-STAT1-KO cells.

In macrophages, no viral replication was detected for the G248V_nsp15_, H250Y_nsp15_ and H250A_nsp15_ mutants (Fig 8A). The K60R_nsp15_ and T66I_nsp15_ mutants showed some growth impairment at 48 h and 72 h (3- to 17-fold reduction vs WT) but this was not significant. In 16HBE cells, most viruses showed very similar growth kinetics as WT virus, with only G248V_nsp15_, H250Y_nsp15_ and H250A_nsp15_ showing a slight (3- to 8-fold) reduction at 48-72 h, which was only significant for H250Y_nsp15_ and H250A_nsp15_ (P ≤ 0.05 at 48 h). In Huh7-STAT1-KO cells, all mutants replicated with the same kinetics as WT virus, the only exception being a slight reduction for the H250A_nsp15_ mutant at 48 h p.i. An intermediate picture was seen in HEL cells (Fig 8A). Here, the H250Y_nsp15_ and H250A_nsp15_ mutants were significantly impaired (7- to 56-fold; P ≤ 0.01) at 24 to 72 h p.i. The G248V_nsp15_ and K60R_nsp15_ mutants showed a modest (4- to 7-fold) reduction, while the T66I_nsp15_ virus was indistinguishable from WT virus.

**Fig 8.**
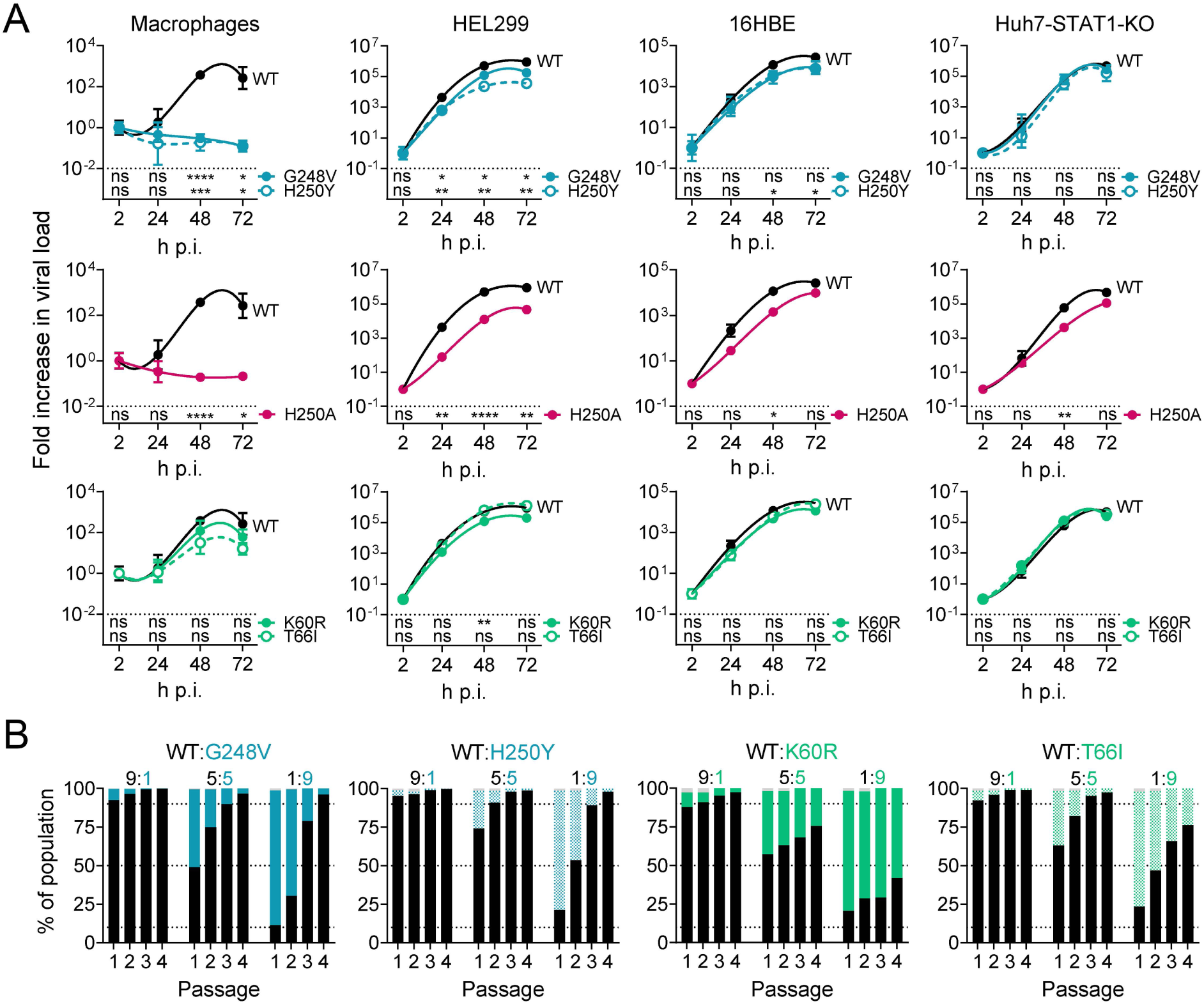
HCoV-229E viruses bearing EPB-113 resistance mutations have impaired viral fitness in IFN responsive cells. (A) Replication kinetics in human macrophages and HEL299, 16HBE and Huh7-STAT1-KO cells, based on RT-qPCR for viral load. Data points are the mean ± SEM (N=3). At each time point, statistical significance is shown for the difference between mutant and WT (multiple unpaired t-tests, with Holm-Šídák’s correction for multiple comparisons). (B) Competitive growth assay. At the start of the experiment, each mutant was mixed with WT virus at a ratio of 9:1, 1:1 or 1:9, then applied to HEL299 cells. After the virus was passaged three more times, Nanopore sequencing was performed on each passage, to determine the percentage of viral genomes that contained the mutant or WT form of nsp15.

#### The mutant viruses are outcompeted by WT virus

To estimate the competitive fitness of the mutants vis-à-vis WT, we co-infected HEL299 cells with three different WT:mutant ratios and then passaged the progeny virus three times; all this virus culturing was done in the absence of compound. Each passage was submitted to Nanopore sequencing to assess the percentage of mutant versus WT sequence in the virus population. This revealed that all nsp15 mutant forms of HCoV-229E were outcompeted by WT virus (Fig 8B). The two catalytic site mutants (G248V_nsp15_ and H250Y_nsp15_) showed the highest fitness cost. Their virus populations almost exclusively consisted of WT after four passages, even if the mutant was 9-fold in excess at the start of the experiment. Also the K60R_nsp15_ and particularly the T66I_nsp15_ virus were readily overtaken by WT virus. Hence, in line with their shifted growth curves in HEL299 cells (see above), the catalytic site mutant viruses proved unable to compete with WT virus. The other two mutations, K60R_nsp15_ and T66I_nsp15_, had less dramatic effect but still rendered the virus less fit in comparison to WT virus.

### Elevated expression of interferon and related players in innate immunity

Parallel to the viral growth experiments, we assessed whether the mutants differed from WT virus in terms of induction of IFN-I and other innate immune factors. The levels of bioactive IFN-α/β protein were determined in human macrophages, HEL299 and 16HBE cells, while all other factors were measured at transcript level and in HEL299 cells only. The data are visualized as heatmaps in Fig 9, while the numeric data and levels of statistical significance are provided in S6 Fig.

**Fig 9.**
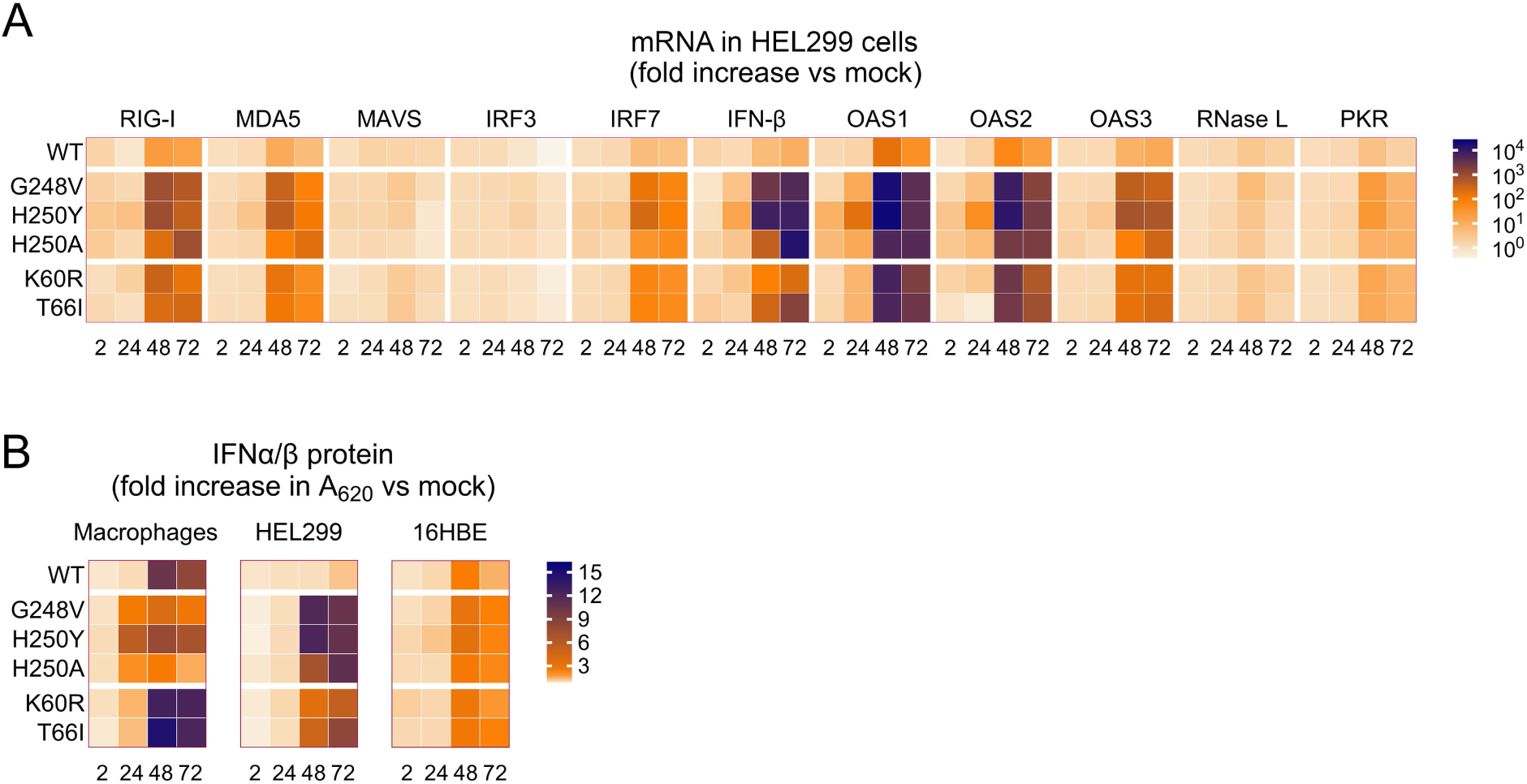
The nsp15-mutant HCoV-229E viruses boost the host cell’s innate immune response. (A) mRNA levels (determined by RT-qPCR) for a range of innate immune factors, in HEL299 cells infected with WT or nsp15-mutant virus. (B) Levels of bioactive IFN-I in the supernatant of human macrophages, HEL299 cells or 16HBE cells infected with WT or mutant virus. For both heatmaps, detailed graphs containing the numeric values and statistical analysis are provided in S6 Fig.

For WT virus (see top row in Fig 9B), the induction of IFN-I was evident at 48 and 72 h but not yet at 24 h p.i. The increases in bioactive IFN-I, compared to mock-infected cells, were up to 10-fold in macrophages, up to 2-fold in 16HBE cells and not seen (i.e. maximum 1.2-fold) in HEL299 cells (Fig 9B). The impact of the nsp15 mutations differed according to time and cell type and, overall, the viruses carrying EndoU catalytic site mutations had a different profile than the K60R_nsp15_ and T66I_nsp15_ mutants. When analyzed at 48 or 72 h p.i., macrophages infected with the K60R_nsp15_ or T66I_nsp15_ virus showed slightly higher levels of IFN-I than cells infected with WT virus. The G248V_nsp15_, H250Y_nsp15_ and H250A_nsp15_ viruses not only showed the opposite profile (i.e. less IFN-I than WT at 48 and 72 h p.i.), but also differed from WT virus in showing an increase in IFN-I at 24 h p.i. Apparently, the levels of IFN-I induced by these mutant viruses within the first 24 h are sufficient to fully suppress their replication (as evident from the growth curves above) which, in turn, prevents a further strong rise in IFN-I at later time points. The early IFN-I induction was not seen in the other two cell types. In 16HBE cells analyzed at 48 h or 72 p.i., the nsp15 mutants showed modest increases in IFN-I versus WT, although the difference reached significance (P ≤ 0.001) for the G248V_nsp15_ and H250Y_nsp15_ viruses. In HEL cells, all nsp15 mutants generated markedly higher IFN-I levels than WT virus; the differences were more significant for the three catalytic site mutants (9-fold; P ≤ 0.0001 at 72 h p.i.) than for K60R_nsp15_ (4-fold; ns) and T66I_nsp15_ (7-fold, P = 0.002).

The panel of innate immune factors, determined at transcript level in HEL299 cells, consisted of (Fig 9A): the pathogen recognition receptors (PRRs) retinoic acid-inducible gene I (RIG-I) and melanoma differentiation-associated protein 5 (MDA5); the signaling factors mitochondrial antiviral-signaling protein (MAVS), interferon regulatory factor 3 (IRF3) and 7 (IRF7); IFN-β; and antiviral effectors 2′-5′-oligoadenylate synthetases (OAS1-3), RNase L and protein kinase R (PKR). All nsp15 mutants generated significantly higher mRNA levels than WT virus for RIG-I, MDA5, IRF7, OAS1-3 and PKR. The increases were only evident at 48 and 72 h p.i., in line with the results obtained for the IFN-I bioassay in HEL299 cells. Particularly abundant expression was seen for OAS-1 and OAS-2: at 48 h p.i., their mRNA levels were 2000- to 25,000-fold increased versus mock, compared to 60- to 160-fold for WT virus, giving high significance (P ≤ 0.001) for each comparison between mutant and WT [see the numeric values (log_10_ scale) and significance levels in S6 Fig). The catalytic site mutants showed higher expression of the innate immune factors than the other two mutants. For instance, for the G248V_nsp15_, H250Y_nsp15_ and H250A_nsp15_ mutants, RIG-I mRNA was up to ∼800-fold increased versus mock, while the K60R_nsp15_ and T66I_nsp15_ mutants showed a maximum increase of ∼300-fold. For MAVS and RNase L, we observed no or only weak induction, regardless of whether the cells were infected with WT or mutant. Finally, we observed that HCoV-229E infection slightly reduces the transcription of IRF3 (3-fold), and that EndoU catalytic site mutations significantly (P ≤ 0.01) revert this effect. This is in line with the finding that PEDV EndoU was reported to degrade the mRNA of IRF3 [62]. We note, however, that our analysis of innate immune factors looked only at their transcript levels, and not at their protein level, status of activation (e.g. from phosphorylation) or downstream activity.

### The mutants are more sensitive to temperature and treatment with IFN-β

Since several studies [63, 64] have shown that the antiviral innate immune response is stronger at 37-38 °C than at 33-35 °C, we assessed the temperature-sensitive growth of the WT and nsp15-mutant HCoV-229E viruses in either IFN responsive (HEL299) or non-responsive (Huh7-STAT1-KO) cells (Fig 10A). Compared to 33-35 °C, all viruses exhibited reduced growth at 37 °C and particularly 38 °C. The latter temperature resulted in marked impairment of viral replication for the H250Y_nsp15_, H250A_nsp15_ and G248V_nsp15_ mutants, however this was only seen in HEL299 cells and not in Huh7-STAT1-KO cells. This aligns with the high innate immunity-inducing phenotype of these mutants (see above). In contrast, for the WT, K60R_nsp15_ and T66I_nsp15_ viruses, the temperature sensitivity was similar in HEL299 and Huh7-STAT1-KO cells, which aligns with the above finding that these viruses induce lower levels of IFN-I than H250Y_nsp15_, H250A_nsp15_ and G248V_nsp15_.

**Fig 10.**
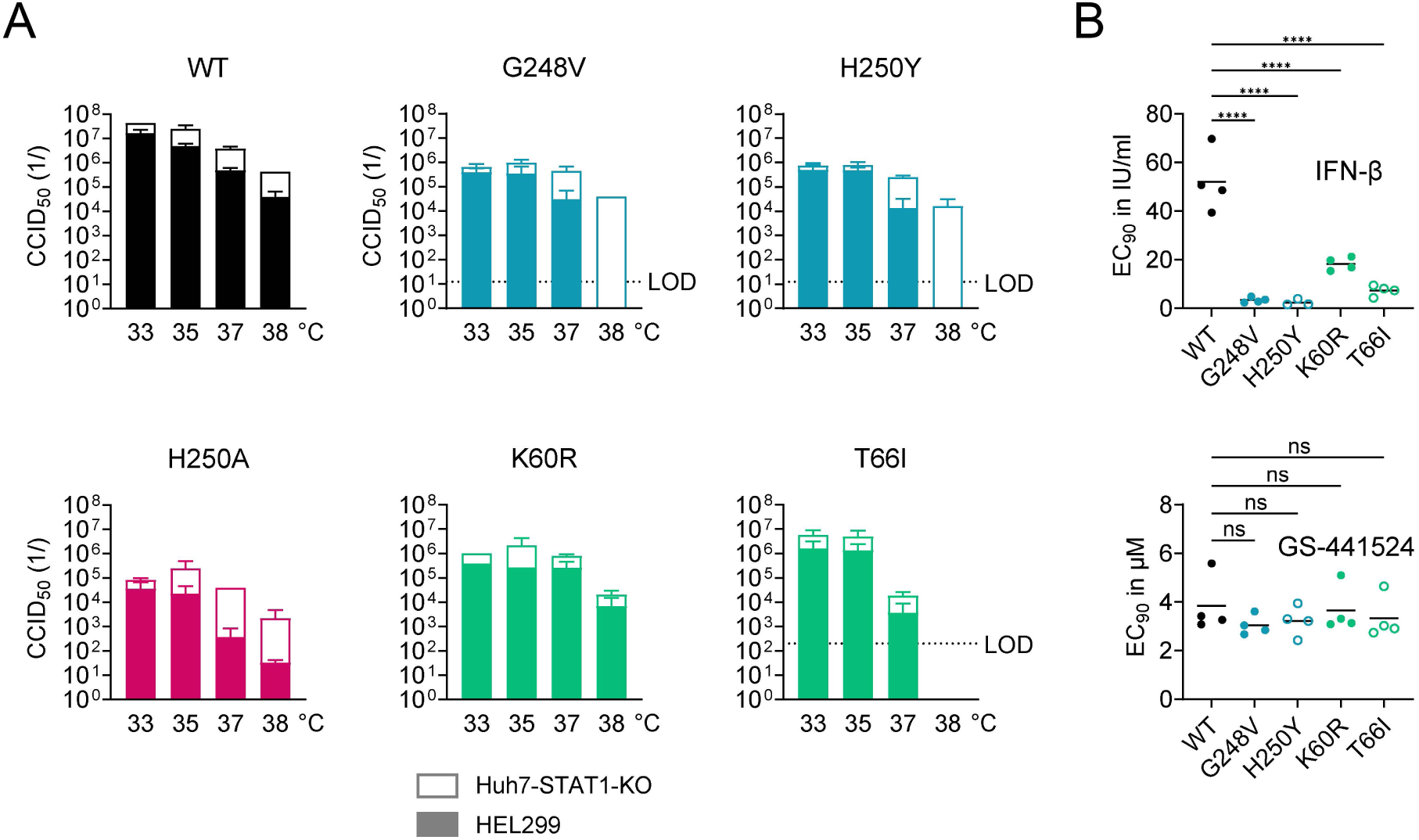
The nsp15 mutations increase the sensitivity to temperature and exogenous IFN-β. (A) Temperature-dependent replication of WT and nsp15-mutant HCoV-229E viruses in IFN responsive (HEL299) or non-responsive (Huh7-STAT1-KO) cells. The graphs show the mean CCID_50_ values of two independent virus titration experiments, conducted at the indicated temperatures. (B) The nsp15 mutations render HCoV-229E more sensitive to the antiviral effect of exogenous IFN-β, as determined by viral load assay in HEL299 cells. The graphs show the individual and mean values from four independent experiments. An ordinary one-way ANOVA with Dunnett’s correction was used to analyze the differences in EC_90_, comparing mutants to WT virus.

Considering that the EndoU-mutants induce higher levels of IFN-I effectors than WT virus, they may also be more responsive to exogenous interferon. This was indeed demonstrated by Kindler et al. [11], for H250A_nsp15_-mutant HCoV-229E evaluated in human MRC5 cells. Hence, we determined the antiviral activity of recombinant human IFN-β in HEL299 cells (Fig 10B), which are the same cell type as MRC5 (i.e. both are human fibroblast cells isolated from normal fetal lung tissue). RT-qPCR for viral load was performed on the supernatants collected at day 3 p.i. Compared to WT virus, all nsp15-mutants exhibited markedly lower EC_90_ values for IFN-β. The G248V_nsp15_ and H250Y_nsp15_ mutants showed a very strong shift of ∼20-fold. The T66I_nsp15_ and K60R_nsp15_ mutants were less affected (= ∼7- and ∼3-fold lower EC_90_ value than WT, respectively), but still showed significantly (P ≤ 0.0001) higher IFN-β sensitivity than WT.

Collectively, the following conclusions can be made from this phenotypic profiling of the nsp15-mutant HCoV-229E viruses which arose under EPB-113 or 5h. Compared to WT virus, all mutants show an altered IFN-I response, but the kinetics, intensity and consequences of this effect depend on the location of the mutation (i.e. in the catalytic site or adjacent RNA binding groove) and cell type. In agreement with two previous studies on EndoU-mutant forms of HCoV-229E and MHV [10, 11], our EndoU catalytic site mutants show early induction of IFN-I, explaining their inability to replicate in macrophages and their severely compromised replication at 38 °C in STAT1-expressing HEL299 cells. The K60R and T66I mutations located in the RNA binding groove cause less dramatic growth impairment in macrophages, but do induce high levels of IFN-I in HEL299 cells. As a consequence all our nsp15 mutants were readily outcompeted by WT virus.

## Discussion

In this study, a detailed mechanistic investigation on the CoV replication inhibitor EPB-113 led to the discovery of a novel antiviral concept targeting the viral nsp15 protein. Additionally, our research highlights the broader importance of nsp15 EndoU for viral replication, which extends beyond its role in evading innate immunity.

Our findings challenge the conventional pharmacological view that disrupting the function of nsp15 in virus-infected cells is best accomplished through inhibition of the EndoU-mediated RNA cleavage reaction. Based on this concept, the search for nsp15 inhibitors has typically relied on an enzymatic FRET assay that is suitable for high-throughput screening. Several hit compounds identified with the nsp15 protein from SARS-CoV [27] or SARS-CoV-2 [28–32] demonstrated favorable IC_50_ values in a FRET or similar enzymatic EndoU assay. However, in all cases reported so far, the compounds showed disappointing inhibitory activity in CoV-infected cells. Since these compounds have diverse chemical structures, it is unlikely that poor cell permeability is the sole issue. Rather, it may be that, unless a compound achieves complete inhibition of EndoU cleavage, viral replication cannot be effectively suppressed. Alternatively, the issue may be methodological: enzymatic EndoU assays may not accurately capture the complex functions of nsp15 in viral replication, while antiviral assays in interferon-deficient cell lines, such as Vero cells, overlook potential IFN induction by an EndoU inhibitor [41].

Our study with EPB-113 introduces a third and previously unrecognized possibility: disrupting the role of nsp15 in virus-infected cells may be achieved with an nsp15 binder that alters the protein’s conformation in a way that enhances EndoU cleavage. The concept of enzyme activation offers several advantages, including the potential to achieve strong efficacy within cells, even if the drug only moderately activates the enzyme [65]. Although the activation by EPB-113 may seem modest based on fluorographs from the FRET assay, this effect was confirmed by HPLC-MS analysis which showed pronounced increase of the 5′-hydroxylated product, the 3′-phosphorylated product and the 2′-3′-cyclic intermediate. EPB-113 seems to induce these effects by binding at the catalytic core of EndoU (since its binding required the His250 and Gly248 residues) and causing a conformational change in the nsp15 hexameric structure, as suggested by the thermal shift assay. The reduction of T_m_ caused by EPB-113 is significant and in contrast with the effect reported for tipiracil [34], a UMP analogue that causes modest inhibition of the EndoU reaction and a slight increase (0.5 °C) in the thermostability of SARS-CoV-2 nsp15. Cryo-EM analysis showed that the nsp15 apoprotein is highly flexible with ‘wobbling’ EndoU domains that are stabilized upon RNA binding [35]. We speculate that EPB-113 might alter conformational flexibility, inducing an RNA-bound state that is catalytically favorable, similar to the mechanism proposed for the EndoU-stimulating effect of Mn^2+^ [66, 67]. Since the positively charged guanidine function (which is fully protonated at physiological pH) is necessary for both the EndoU stimulation and strong antiviral activity, EPB-113 might perhaps enhance the interaction between the RNA ligand and the negatively charged surface of EndoU [36]. The idea that EPB-113 acts as a catalytic activator of EndoU aligns with its resistance profile, as the virus escaped the compound by losing this enzymatic activity. An analysis of the physiologic substrates of MHV EndoU suggested that its cleavage of specific motifs in viral positive-sense RNA occurs in a regulated manner, to partially inhibit negative-strand RNA synthesis and prevent accumulation of viral dsRNA [9]. More recently, SARS-CoV-2 EndoU was proposed to regulate the process of recombination for discontinuous synthesis of the subgenomic viral mRNAs and production of defective viral genomes [25]. We hence hypothesize that EPB-113 acts by tilting the balanced EndoU activity towards a level that is not compatible with efficient viral RNA synthesis.

As the first nsp15 binder with strong anti-CoV activity in infected cells, EPB-113 represents a unique tool for unravelling the functions of EndoU beyond its role in immune evasion. The potent anti-HCoV-229E activity in STAT1-deficient cells means that EPB-113 mainly uses an IFN-independent mechanism to suppress viral replication. Intriguingly, in HEL299 cells, we observed a subtle increase in IFN-I, but only when EPB-113 was added at an intermediate concentration (∼1 µM) and at 24 h p.i. This limited boosting of IFN-I provides further support that EPB-113 does not act by EndoU inhibition. To explain why EPB-113 induces IFN-I in certain circumstances, analysis would be needed of the different forms of viral RNA in EPB-113-treated cells, combined with detailed understanding of how RNA synthesis and immune evasion is regulated by EndoU cleavage at specific motifs in viral negative- or positive-sense RNA [9, 12, 25].

The resistance profile of EPB-113 is another interesting feature, since the compound selects for EndoU-deficient mutant viruses having reduced fitness in IFN-competent cells. Our findings confirm that, at least for HCoV-229E, having a functional EndoU offers a clear advantage to the virus. At first sight, this seems contradictory to reports of efficient replication of EndoU-deficient CoVs in various cell models [23, 68]. However, these studies typically use immortalized cell lines, many of which have defects in the IFN-I pathway. Consistent with a previous study [11], we confirm that EndoU-deficient HCoV-229E viruses (bearing nsp15 mutations G248V, H250Y or H250A) trigger an early IFN-I response in human macrophages, rendering them unable to replicate in these cells. While the mutants replicate efficiently in human lung fibroblasts, virus competition experiments reveal that they are easily outcompeted by WT virus. The vulnerability of the mutants stems from their ability to excessively stimulate the innate immune response, activating several layers of defense, including IFN-I, PRRs, signaling factors and antiviral effectors. This amplifies the suppressive effect of IFN-I on the replication of the nsp15 mutants, leading to their elimination when WT virus is also present.

In a recent study, nsp15 was shown to downregulate the cGAS-STING innate immunity pathway, with the HCoV-229E protein having apparently stronger effect than the SARS-CoV-2 analogue [26]. Hence, CoVs may show subtle differences in their reliance on EndoU. Mutations G248V_nsp15_, H250Y_nsp15_ and H250A_nsp15_, are viable for HCoV-229E, while reverse-engineering these changes in SARS-CoV-2 did not yield viable viruses. However, we successfully rescued two reverse-engineered SARS-CoV-2 mutants with substitution K60R_nsp15_ or V66I_nsp15_ in the RNA binding groove, adjacent to the catalytic site. Using these mutants, we confirmed that nsp15 is also the target for EPB-113 in SARS-CoV-2. Future development of EPB-113 will require structural optimization, to achieve sufficient anti-SARS-CoV-2 activity.

In conclusion, while the many unknowns regarding the function of nsp15 in CoV replication present a challenge for antiviral drug development, our study demonstrates that nsp15 is a valid target for inhibiting CoV replication. We provide evidence that strong antiviral activity can be achieved with a compound, like EPB-113, which alters the conformation of nsp15 and enhances its EndoU activity up to a level that is not tolerated by the replicating virus. Escaping such a compound requires that the virus loses EndoU-activity, but comes at a serious fitness cost. This makes the concept of nsp15 binders highly attractive for antiviral therapy.

## Acknowledgements

We wish to thank L. Delarbre, B. Ramos, N. Willems and S. Wouters for excellent technical assistance and V. Thiel, B. Canard, P. Hoet, P. J. Bredenbeek, D. Jochmans, X. Saelens and P.-Y. Shi for the kind gift of viruses, plasmids or cells. Images were produced using Graphpad Prism 10.3.1 (GraphPad Software, Boston, Massachusetts USA), the UCSF Chimera package [69] from the Resource for Biocomputing, Visualization, and Informatics at the University of California, San Francisco (supported by NIH P41-GM103311); ESPript [70], SynergyFinder+ [55] and Marvin (https://www.chemaxon.com).

## S1 Appendix Chemical synthesis and analysis of EPB-113 and its two structural analogues EPB-102 and JMLv-061

### S1 Appendix: Chemistry

#### Chemical synthesis. General methods

Commercially available reagents and solvents were used without further purification unless stated otherwise. Preparative normal phase chromatography was performed on a CombiFlash Rf 150 (Teledyne Isco) with pre-packed RediSep Rf silica gel cartridges. Thin-layer chromatography was performed with aluminum-backed sheets with silica gel 60 F_254_ (Merck, ref 1.05554), and spots were visualized with UV light and 1% aqueous solution of KMnO_4_. Melting points were determined in open capillary tubes with a MFB 595010M Gallenkamp. 400 MHz ^1^H, 100.6 MHz ^13^C and 376.5 MHz ^19^F NMR spectra were recorded on a Varian Mercury 400 or on a Bruker 400 Avance III spectrometers. The chemical shifts are reported in ppm (*δ* scale) relative to internal tetramethylsilane, and coupling constants are reported in Hertz (Hz). Assignments given for the NMR spectra of the new compounds have been carried out on the basis of COSY ^1^H/^13^C (gHSQC sequence) experiments. IR spectra were run on a Perkin-Elmer Spectrum RX I spectrophotometer. Absorption values are expressed as wave-numbers (cm^−1^); only significant absorption bands are given. High-resolution mass spectrometry (HRMS) analyses were performed with an LC/MSD TOF Agilent Technologies spectrometer. The elemental analyses were carried out in a Flash 1112 series Thermofinnigan elemental microanalyzator (A5) to determine C, H, N and S. The structure of all new compounds was confirmed by elemental analysis and/or accurate mass measurement, IR, ^1^H NMR, ^13^C NMR and ^19^F NMR. The analytical samples of all the new compounds, which were subjected to antiviral evaluation, possessed purity ≥ 95% as evidenced by their elemental analyses and/or their HPLC/MS. HPLC/MS for EPB-113 was determined with a HPLC Agilent 1260 Infinity II LC/MSD coupled to a photodiode array and mass spectrometer. 5 µL of sample 0.5 mg/mL in methanol:acetonitrile were injected, using a Agilent Poroshell 120 EC-C18, 2.7 µm, 50 mm x 4.6 mm column at 40 °C. The mobile phase was a mixture of A = water with 0.05% formic acid and B = acetonitrile with 0.05% formic acid, with the method described as follows: flow 0.6 mL/min, from 95% A - 5% B to 100% B in 3 min, 100% B 3 min, from 100% B to 95% A - 5% B in 1 min, 95% A – 5% B 3 min. Purity is given as % of absorbance at 254 nm. Data from mass spectra were analyzed by electrospray ionization in positive and negative between 100 and 1000 Da.

#### Synthesis of 1-(3-aminophenyl)-3-(3-(pentafluoro-λ^6^-sulfanyl)phenyl)urea, EPB-102

##### Step 1: Synthesis of 1-(3-nitrophenyl)-3-(3-(pentafluoro-λ^6^-sulfanyl)phenyl)urea

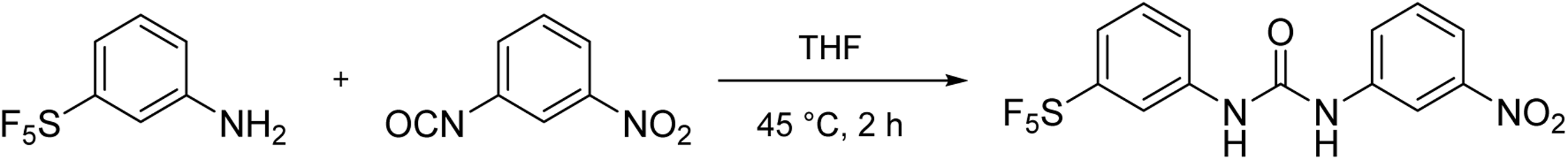

3-Nitrophenylisocyanate (500 mg, 3.05 mmol) was dissolved in THF (25 mL) and 3-pentafluorosulfanyl aniline (668 mg, 3.05 mmol) was added. The reaction mixture was stirred at 45 °C for 2 hours. The solvent was removed under reduced pressure to afford a yellow solid (1.10 g) that was crystallized in EtOAc to give pure 1-(3-nitrophenyl)-3-(3-(pentafluoro-λ^6^-sulfanyl)phenyl)urea (1.10 g, 94% yield) as a yellow crystalline solid.

##### Step 2: Synthesis of EPB-102

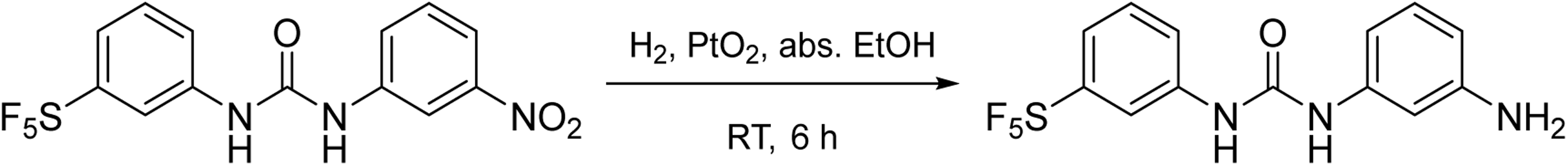

A suspension of 1-(3-nitrophenyl)-3-(3-(pentafluoro-λ^6^-sulfanyl)phenyl)urea (600 mg, 1.56 mmol) and PtO_2_ hydrate (53 mg) in abs. ethanol (45 mL) was hydrogenated (1 atm) at room temperature for 6 hours. The suspension was then filtered, and the solvent was evaporated under vacuum to afford **EPB-102** (533 mg, 97% yield) as a white solid. The analytical sample was obtained by crystallization from EtOAc, mp (EtOAc) 193–194 °C. IR (ATR) ν: 643, 651, 682, 697, 789, 824, 845, 877, 913, 1167, 1229, 1313, 1420, 1465, 1485, 1566, 1590, 1651, 3092, 3294, 3389, 3489 cm^-1^. ^1^H-NMR (400 MHz, DMSO-*d*_6_) δ: 5.05 (s, 2 H, NH_2_), 6.21 (m, 1 H, 6-H or 4-H), 6.52 (m, 1 H, 4-H or 6-H), 6.84 (t, *J* = 2.0 Hz, 1 H, 2-H), 6.90 (t, *J* = 8.0 Hz, 1 H, 5-H), 7.40-7.55 (complex signal, 3 H, 4’-H, 5’-H, 6’-H), 8.28 (t, *J* = 2.0 Hz, 1 H, 2’-H), 8.51 (s, 1 H) and 9.03 (s, 1 H) (2 NH). ^13^C-NMR (100.6 MHz, DMSO-*d*_6_) δ: 104.1 (CH, C2), 106.3 (CH, C4 or C6), 108.4 (CH, C6 or C4), 114.6 (quint, ^3^*J*_CF_ = 5.0 Hz, CH, C2’), 118.5 (quint, ^3^*J*_CF_ = 3.5 Hz, CH, C4’), 121.5 (CH, C6’), 129.0 (CH, C5), 129.6 (CH, C5’), 139.8 (C, C1), 140.7 (C, C1’), 149.2 (C, C3), 152.3 (C, CO), 153.3 (quint, ^2^*J*_CF_ = 15.1 Hz, C, C3’). ^19^F-NMR (376.5 MHz, DMSO-*d*_6_) δ: 63.6 (d, *J* = 150.6 Hz, 4 F, S<u>F</u>_4_F), 87.8 (quint, *J* = 150.6 Hz, 1 F, SF_4_<u>F</u>). HRMS-ESI^-^ *m/z* [M-H]^-^ calculated for [C_13_H_12_F_5_N_3_OS-H]^-^: 352.0548. Found: 352.0554. Elemental analysis calculated for C_13_H_12_F_5_N_3_OS: C 44.19%, H 3.42%, N 11.89%, S 9.07%. Found: C 44.24%, H 3.44%, N 11.63%, S 8.78%.

#### Synthesis of 1-(3-guanidinophenyl)-3-(3-(pentafluoro-λ^6^-sulfanyl)-phenyl)urea hydrochloride, EPB-113

##### Step 1: Synthesis of 2-(3-nitrophenyl)-1,3-di-Boc-guanidine

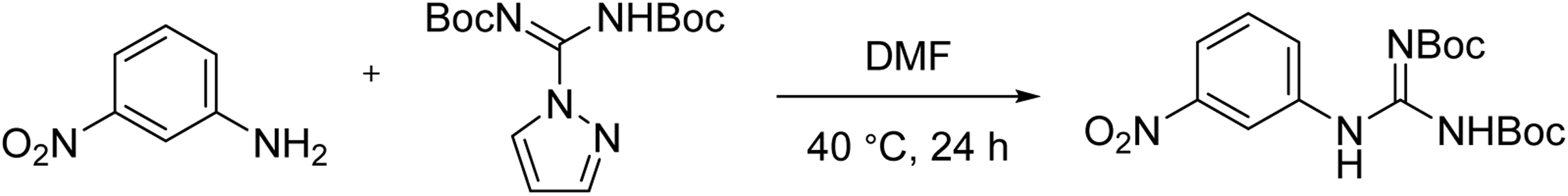

To a stirred solution of 3-nitroaniline (1 g, 7.24 mmol) in DMF (10 mL) was added bis-Boc-pyrazolocarboxamidine (2.25 g, 7.25 mmol) and the reaction mixture was stirred at 40 °C for 24 hours. Water (15 mL) and EtOAc (15 mL) were added, and the phases were separated. The organic layer was further washed with water, dried over anhydrous Na_2_SO_4_ and filtered. Evaporation of the organics provided a yellowish gum (2.46 g), which was purified by column chromatography in silica gel (hexane/EtOAc mixtures). Fractions containing the desired product were collected and concentrated under vacuum to afford 2-(3-nitrophenyl)-1,3-di-Boc-guanidine (1.0 g, 36% yield) as a yellowish solid. The spectroscopic data matched with those previously described in the bibliography.^1^ HRMS-ESI^+^ *m/z* [M+H]^+^ calculated for [C_17_H_24_N_4_O_6_+H]^+^: 381.1769. Found: 381.1769.

##### Step 2: Obtention of 2-(3-aminophenyl)-1,3-di-Boc-guanidine

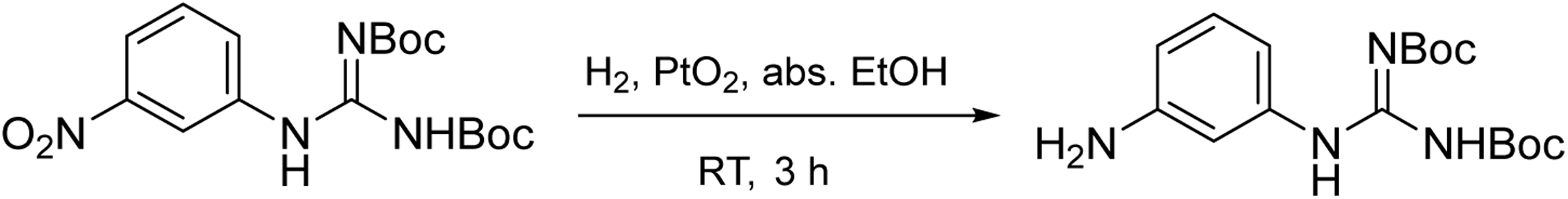

A suspension of 2-(3-nitrophenyl)-1,3-di-Boc-guanidine (1 g, 2.63 mmol) and PtO_2_ hydrate (100 mg) in abs. ethanol (100 mL) was hydrogenated (1 atm) at room temperature for 3 hours. The suspension was then filtered, and the solvent was evaporated under vacuum to afford 2-(3-aminophenyl)-1,3-di-Boc-guanidine (0.91 g, quantitative yield) as a white foamy solid. The spectroscopic data matched with those previously described in the bibliography.^2^

##### Step 3: Synthesis of 1-(1,3-di-Boc-3-guanidinophenyl)-3-(3-(pentafluoro-λ^6^-sulfanyl)phenyl)urea

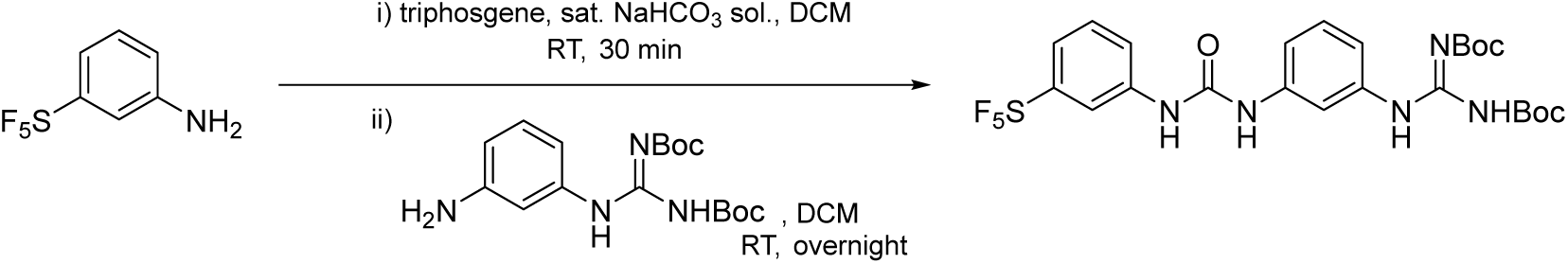

i. 3-pentafluorosulfanyl aniline (80 mg, 0.36 mmol) was added to a stirring biphasic mixture of DCM (5 mL) and saturated aqueous NaHCO_3_ solution (5 mL). Triphosgene (40 mg, 0.14 mmol) was then slowly added and the reaction mixture was stirred at room temperature for 30 minutes. The phases were separated and the organic layer was washed with brine, dried over anhydrous Na_2_SO_4_, filtered and half concentrated at room temperature to give 3-(pentafluoro-λ^6^-sulfanyl)phenylisocyanate in DCM solution that was used in the next step without further purification.
ii. 2-(3-aminophenyl)-1,3-di-Boc-guanidine (139 mg, 0.40 mmol) dissolved in DCM (2 mL) was added to the previously obtained isocyanate in DCM (ca. 6 mL). The reaction mixture was stirred at room temperature overnight. Evaporation under vacuum of the solvent provided an orange solid (350 mg) which was purified by column chromatography in silica gel (hexane/EtOAc mixtures). Fractions containing the desired product were collected and concentrated under vacuum to give 1-(1,3-di-Boc-3-guanidinophenyl)-3-(3-(pentafluoro-λ^6^-sulfanyl)phenyl)urea (60 mg, 29% overall yield) as a yellowish solid. The analytical sample was obtained by washing with Et_2_O. ^1^H-NMR (400 MHz, DMSO-*d*_6_) δ: 1.41 (s, 9 H, Boc), 1.51 (s, 9 H, Boc), 7.15-7.35 (complex signal, 3 H, 2-H, 4-H, 6-H), 7.41-7.59 (complex signal, 3 H, 4’-H, 5-H, 6’-H), 7.64 (s, 1 H, 5’-H), 8.27 (m, 1 H, 2’-H), 8.93 (s, 1 H) and 9.17 (s, 1 H) (2 NH urea), 9.98 (s, 1 H) and 11.35 (s, 1 H) (2 NH guanidine).

##### Step 4: Synthesis of EPB-113 (as its hydrochloride salt)

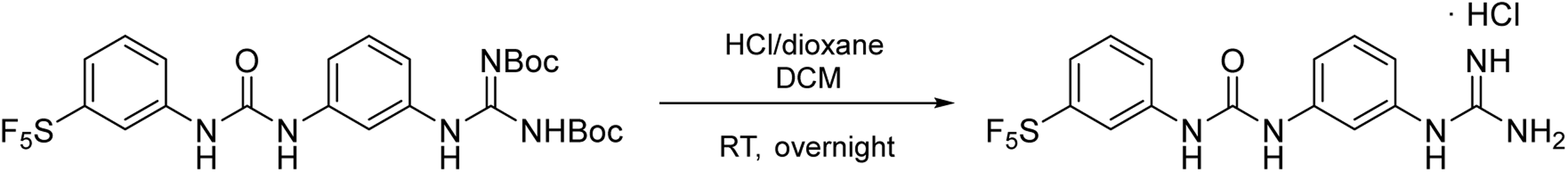

To a stirred suspension of 1-(1,3-di-Boc-3-guanidinophenyl)-3-(3-(pentafluoro-λ^6^-sulfanyl)phenyl)urea (32 mg, 0.05 mmol) in DCM (1 mL) was added a 4 M solution of HCl in 1,4-dioxane (4 mL, 16 mmol) and the mixture was stirred at room temperature overnight. Evaporation of the solvents provided a beige solid, which was washed with Et_2_O and pentane to give **EPB-113** as its hydrochloride salt (19 mg, 88% yield), mp 219–220 °C. IR (ATR) ν: 754, 782, 812, 835, 857, 916, 1098, 1239, 1296, 1406, 1428, 1484, 1496, 1542, 1587, 1598, 1673, 2966, 3184, 3235, 3416 cm^-1^. ^1^H-NMR (400 MHz, DMSO-*d*_6_) δ: 6.86 (m, 1 H, 4-H), 7.27 (m, 1 H, 6-H), 7.35 (t, *J* = 8.0 Hz, 1 H, 5-H), 7.38 (broad s, 4 H, 2 N<u>H</u>_2_ guanidine), 7.46-7.55 (complex signal, 4 H, 2-H, 4’-H, 5’-H, 6’-H), 8.27 (m, 1 H, 2’-H), 9.31 (s, 1 H) and 9.53 (s, 1 H) (2 NH urea), 9.70 (s, 1 H, 3-NH). ^13^C-NMR (100.6 MHz, CD_3_OD) δ: 117.2 (CH, C2), 117.4 (quint, ^3^*J*_CF_ = 5.0 Hz, CH, C2’), 119.4 (CH, C6), 120.5 (CH, C4), 120.9 (quint, ^3^*J*_CF_ = 5.0 Hz, CH, C4’), 123.1 (CH, C6’), 130.3 (CH, C5’), 131.4 (CH, C5), 136.6 (C, C1), 141.3 (C, C1’), 142.0 (C, C3), 154.8 (C, CO), 155.4 (quint, ^2^*J*_CF_ = 17.1 Hz, C, C3’), 158.0 (C, CN guanidine). ^19^F-NMR (376.5 MHz, DMSO-*d*_6_) δ: 63.6 (d, *J* = 150.6 Hz, 4 F, S<u>F</u>_4_F), 87.7 (quint, *J* = 150.6 Hz, 1 F, SF_4_<u>F</u>). HRMS-ESI^+^ *m/z* [M+H]^+^ calculated for [C_14_H_14_F_5_N_5_OS+H]^+^: 396.0912. Found: 396.0916. HPLC (254 nm): t_R_ = 3.75 min (98.4%). Elemental analysis calculated for C_14_H_14_F_5_N_5_OSꞏHClꞏ0.4H_2_O: C 38.30%, H 3.63%, N 15.95%. Found: C 38.51%, H 3.88%, N 16.21%.

#### Synthesis of 1-(3-guanidinophenyl)-3-(2-(trifluoromethoxy)phenyl)urea dihydrochloride, JMLV-061

##### Step 1: Synthesis of 2-(3-nitrophenyl)-1,3-di-Boc-guanidine

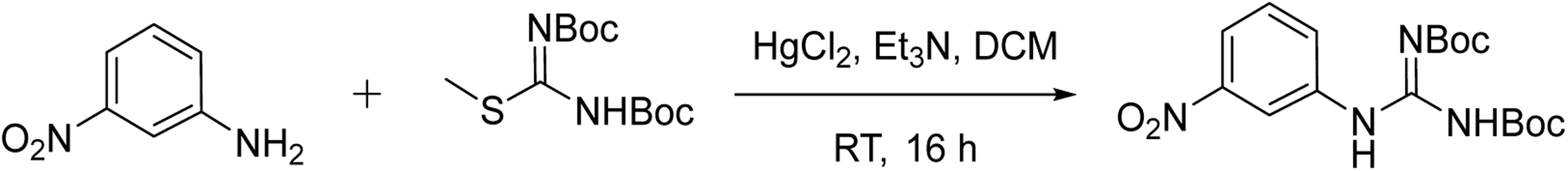

To a solution of 3-nitroaniline (1.50 g, 10.86 mmol) and triethylamine (4.54 mL, 32.58 mmol) in anhydrous DCM (40 mL) was added *N,N’*-bis(tert-butoxycarbonyl)-S-methylisothiourea (3.31 g, 11.40 mmol) and HgCl_2_ (3.24 g, 11.95 mmol) and the mixture was stirred at RT for 16 hours. The mixture was filtered through a pad of Celite® using DCM as eluting agent. Then, the solution was washed with water (2 x 80 mL). The organic layer was dried over anhydrous Na_2_SO_4_ and filtered. Solvents were concentrated under vacuum and the resulting crude was purified by column chromatography in silica gel (hexane/EtOAc mixtures). Fractions containing the desired product were collected and concentrated under vacuum to afford 2-(3-nitrophenyl)-1,3-di-Boc-guanidine as a pale-yellow solid (3.88 g, 94% yield). The spectroscopic data matched with those previously described in the bibliography.^1^ HRMS-ESI^+^ *m/z* [M+H]^+^ calculated for [C_17_H_24_N_4_O_6_+H]^+^: 381.1769. Found: 381.1769.

##### Step 2: Obtention of 2-(3-aminophenyl)-1,3-di-Boc-guanidine

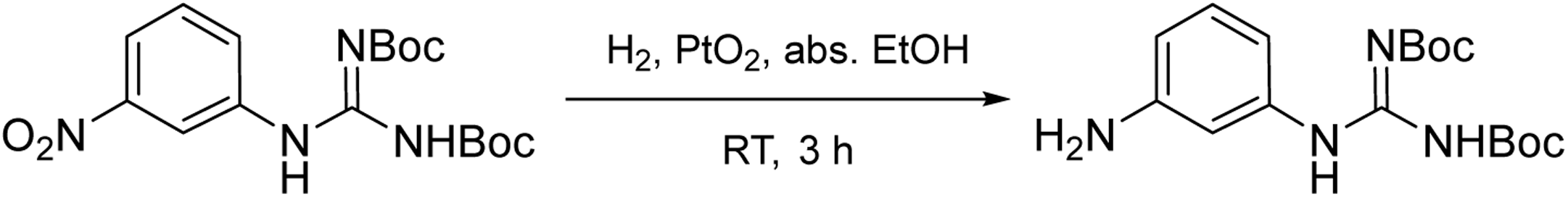

A suspension of 2-(3-nitrophenyl)-1,3-di-Boc-guanidine (2.28 g, 5.99 mmol) and PtO_2_ hydrate (250 mg) in abs. ethanol (50 mL) was hydrogenated at room temperature and atmospheric pressure for 3 hours. Anhydrous Na_2_SO_4_ and Celite^®^ were added and the mixture was stirred for 2 minutes. Then, the suspension was filtered and the solvent was evaporated under vacuum. The resulting crude was purified by column chromatography in silica gel (hexane/EtOAc mixtures). Fractions containing the desired product were collected and concentrated under vacuum to afford 2-(3-aminophenyl)-1,3-di-Boc-guanidine (1.75 g, 83% yield) as a white solid. The spectroscopic data matched with those previously described in the bibliography.^2^

##### Step 3: Synthesis of JMLV-061 (as its hydrochloride)

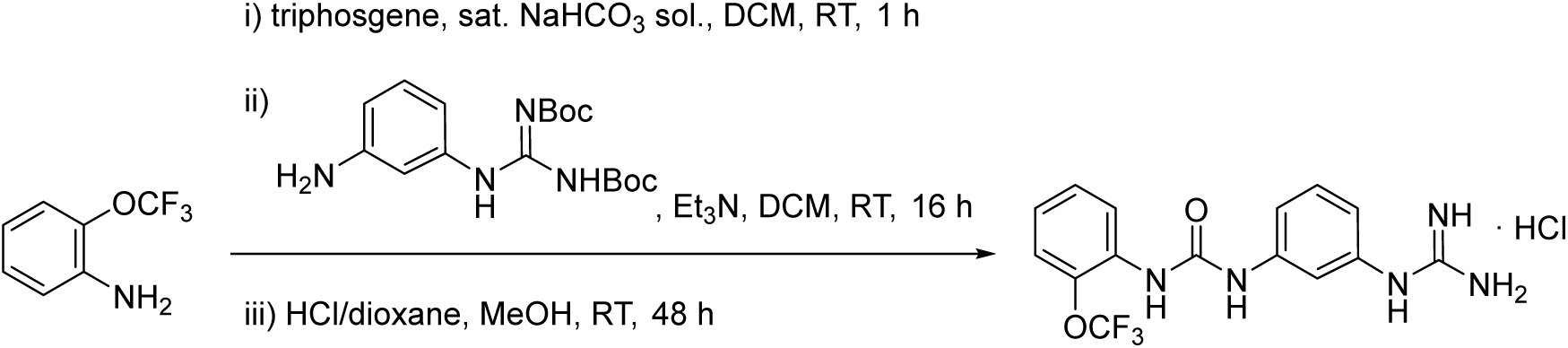

i. To a stirring biphasic mixture of DCM (3 mL), saturated aqueous NaHCO_3_ solution (3 mL) and triphosgene (101 mg, 0.34 mmol) was added 2-(trifluoromethoxy)aniline (93 µL, 120 mg, 0.68 mmol) portionwise. The mixture was then stirred at room temperature for 1 hour. The phases were separated and the organic layer was washed with brine, dried over anhydrous Na_2_SO_4_ and filtered. Solvent was concentrated until 1 mL of DCM was left to give 2-(trifluoromethoxy)phenylisocyanate in solution that was used in the next step without further purification.
ii. 2-(3-aminophenyl)-1,3-di-Boc-guanidine (80 mg, 0.23 mmol) and triethylamine (128 µL, 93 mg, 0.92 mmol) dissolved in DCM (2 mL) was added to the previous isocyanate solution and the mixture was stirred at room temperature for 16 hours. Saturated aqueous NaHCO_3_ solution (15 mL) followed by DCM (10 mL) were added to the mixture and layers were separated. The organic layer was washed again with saturated aqueous NaHCO_3_ solution (15 mL). Then, the organic layer was separated, dried over anhydrous Na_2_SO_4_, filtered and solvents were concentrated under vacuum to afford 1-(1,3-di-Boc-3-guanidinophenyl)-3-(2-(trifluoromethoxy)phenyl)urea that was used in the next step without further purification.
iii. A 4 M solution of HCl/1,4-dioxane (2 mL) and MeOH (0.5 mL) were added to the previously obtained crude and the mixture was stirred at room temperature for 48 hours. The solvents were concentrated under vacuum and the resulting crude was purified by column chromatography in silica gel (DCM/MeOH mixtures). Fractions containing the desired product were collected and concentrated under vacuum to afford **JMLv-061** in its dihydrochloride salt as a beige solid (39 mg, 48% yield), mp 218–219 °C. IR (ATR) ν: 624, 691, 758, 871, 1006, 1059, 1180, 1215, 1314, 1406, 1454, 1490, 1540, 1598, 1674, 2421, 3200, 3330 cm^−1^. ^1^H-NMR (400 MHz, CD_3_OD) δ: 6.95 (ddd, *J* = 8.0 Hz, *J’* = 2.1 Hz, *J’’* = 1.0 Hz, 1 H, 4-H), 7.11 (ddd, *J* = 8.1 Hz, *J’* = 7.5 Hz, *J’’* = 1.6 Hz, 1 H, 4’-H), 7.29 (m, 1 H, 6-H), 7.30-7.34 (complex signal, 2 H, 3’-H, 5’-H), 7.40 (t, *J* = 8.0 Hz, 1 H, 5-H), 7.64 (t, *J* = 2.1 Hz, 1 H, 2-H), 8.20 (dd, *J* = 8.8 Hz, *J’* = 1.6 Hz, 1 H, 6’-H). ^13^C-NMR (100.6 MHz, CD_3_OD) δ: 116.8 (CH, C2), 119.0 (CH, C6), 120.4 (CH, C4), 122.11 (q, *^1^J_CF_* = 256.5 Hz, C, OCF_3_), 122.13 (CH, C3’), 123.1 (CH, C6’), 124.4 (CH, C4’), 128.7 (CH, C5’), 131.5 (CH, C5), 133.1 (C, C1’), 136.7 (C, C1), 139.9 (C, C2’), 142.1 (C, C3), 154.6 (C, CO), 158.0 (C, CN guanidine). HRMS-ESI^+^ *m/z* [M+H]^+^ calculated for [C_15_H_14_F_3_N_5_O_2_+H]^+^: 354.1172. Found: 354.1174. Elemental analysis calculated for C_15_H_14_F_3_N_5_O_2_ꞏHClꞏ0.6DCM: C 42.51%, H 3.70%, N 15.89%. Found: C 42.38%, H 3.90%, N 15.70%.

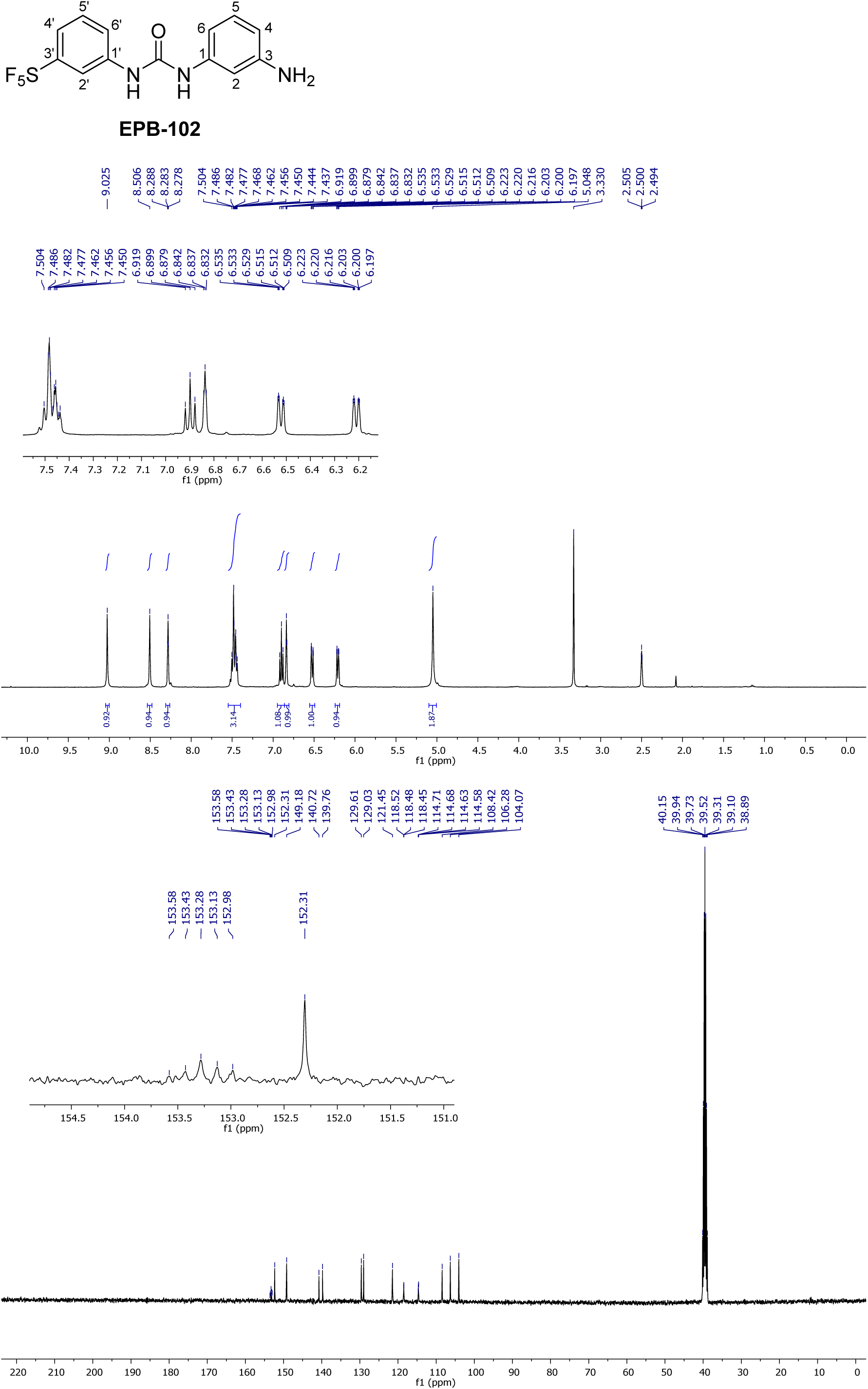

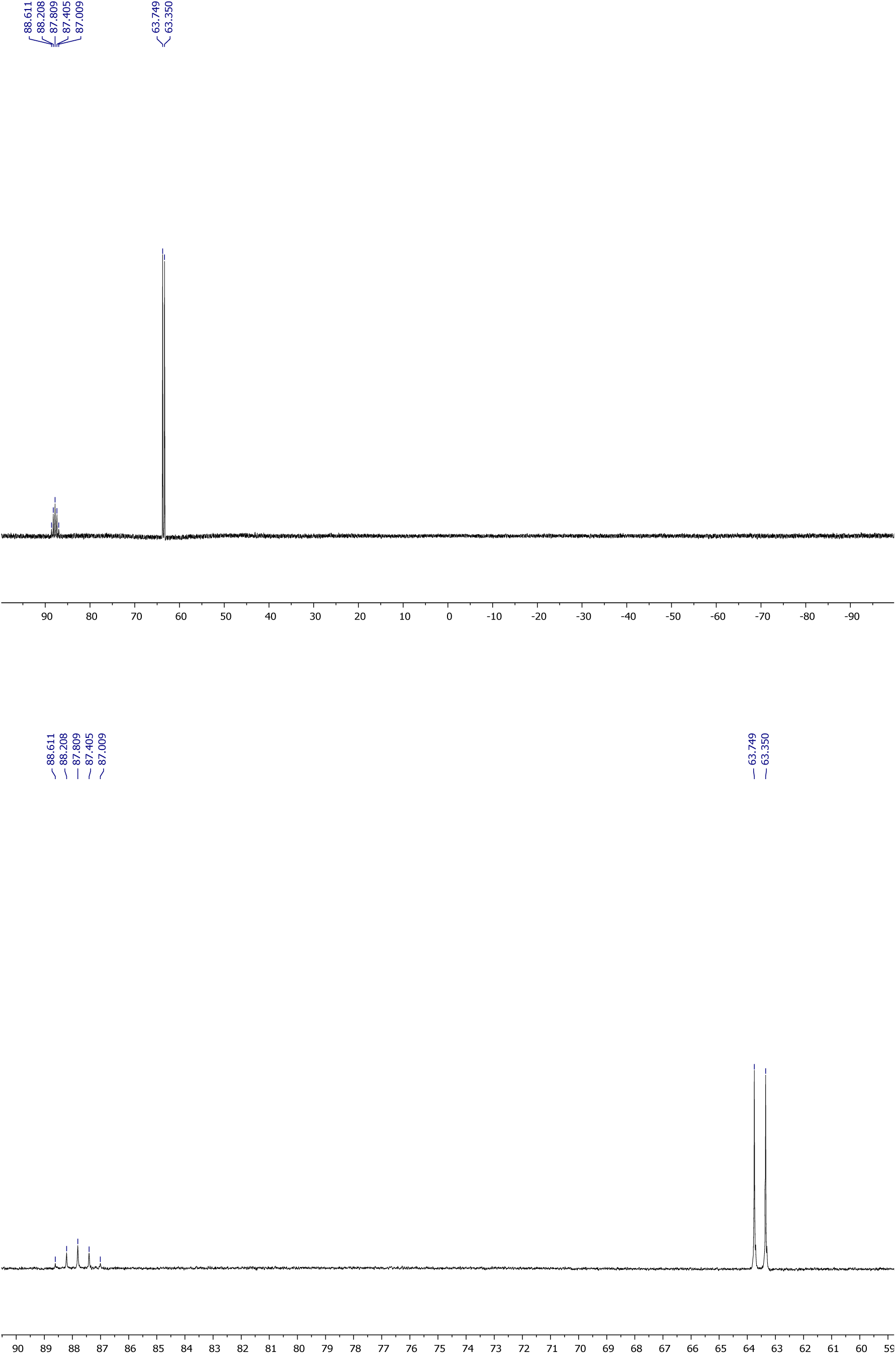

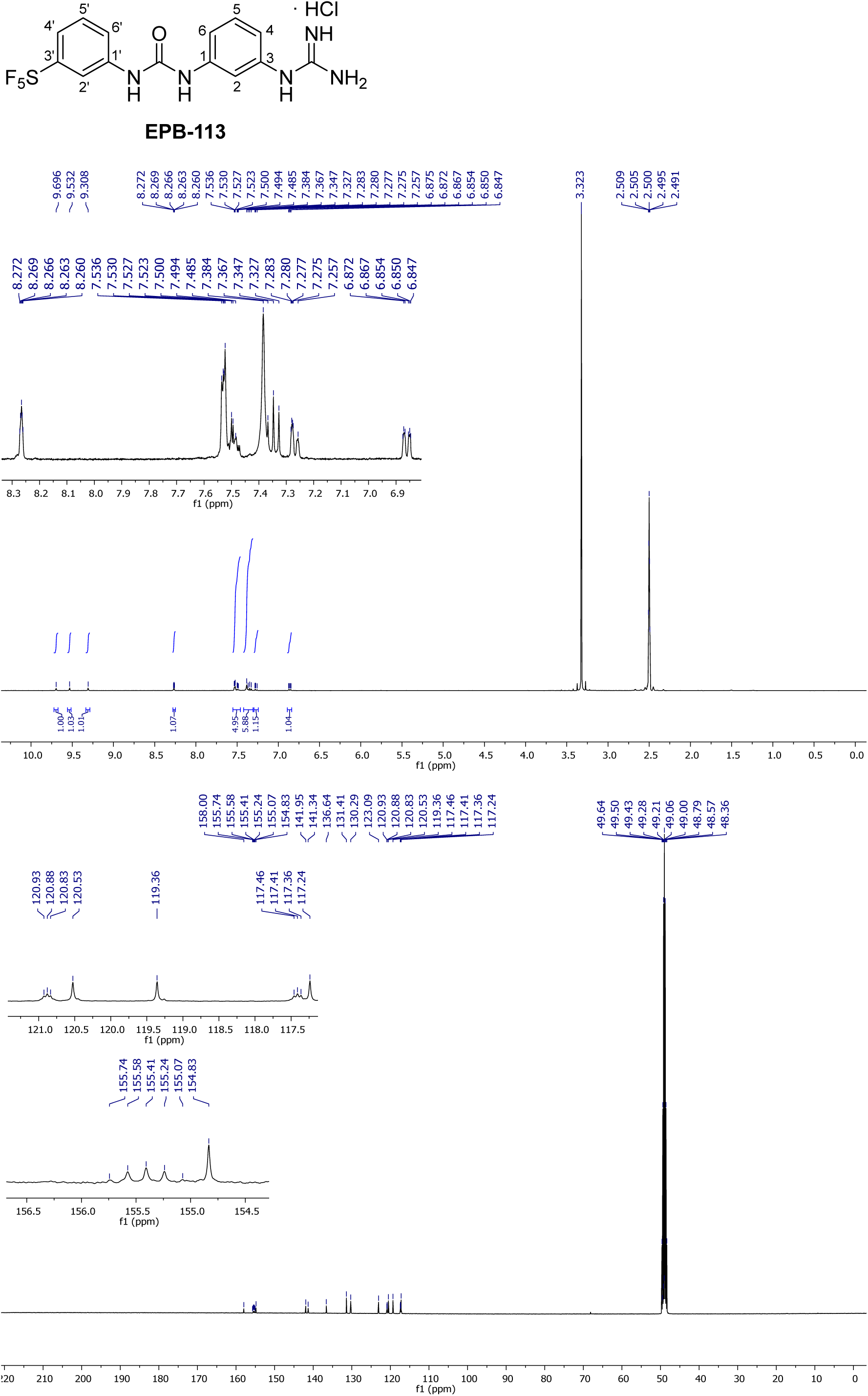

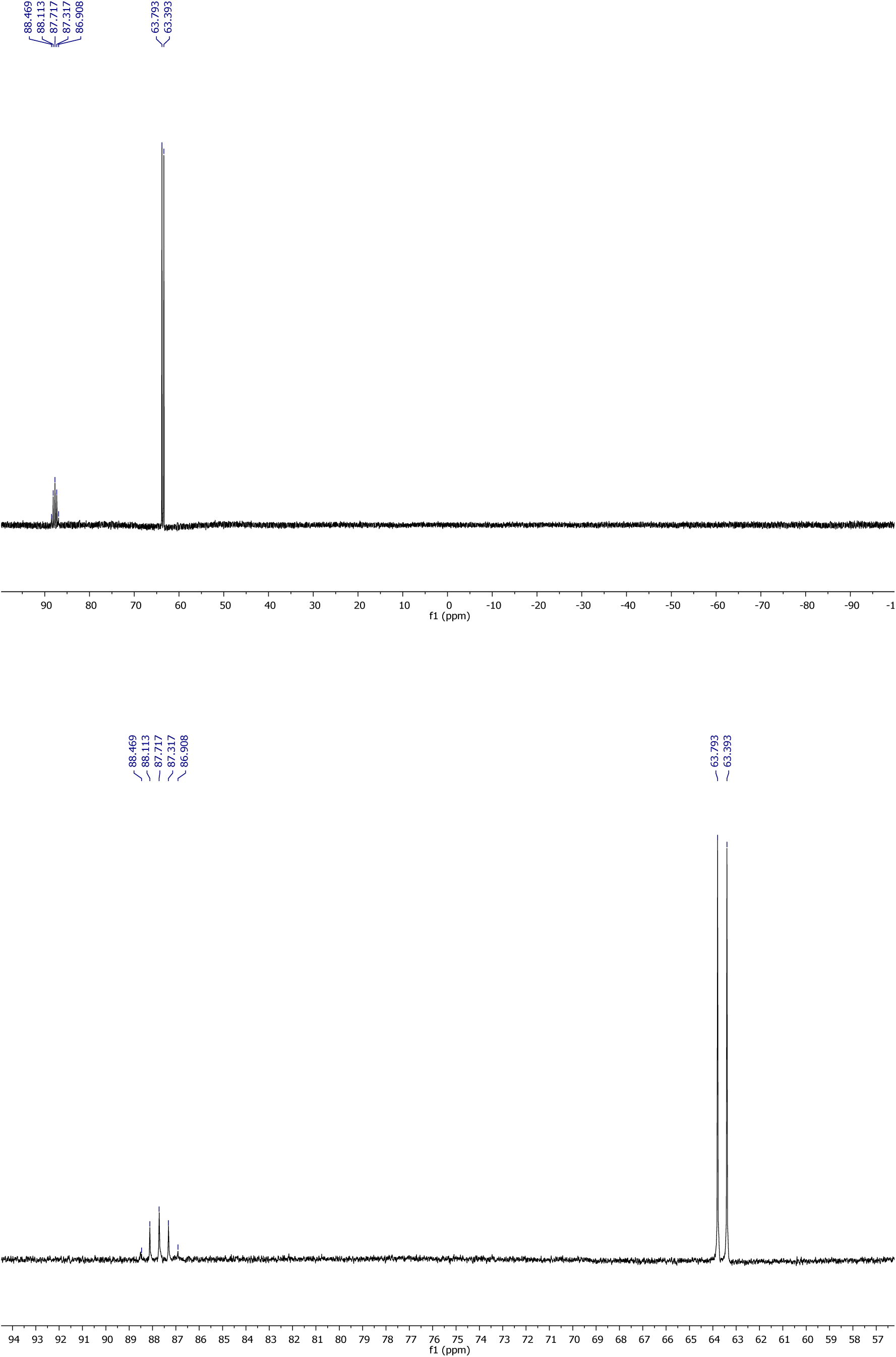

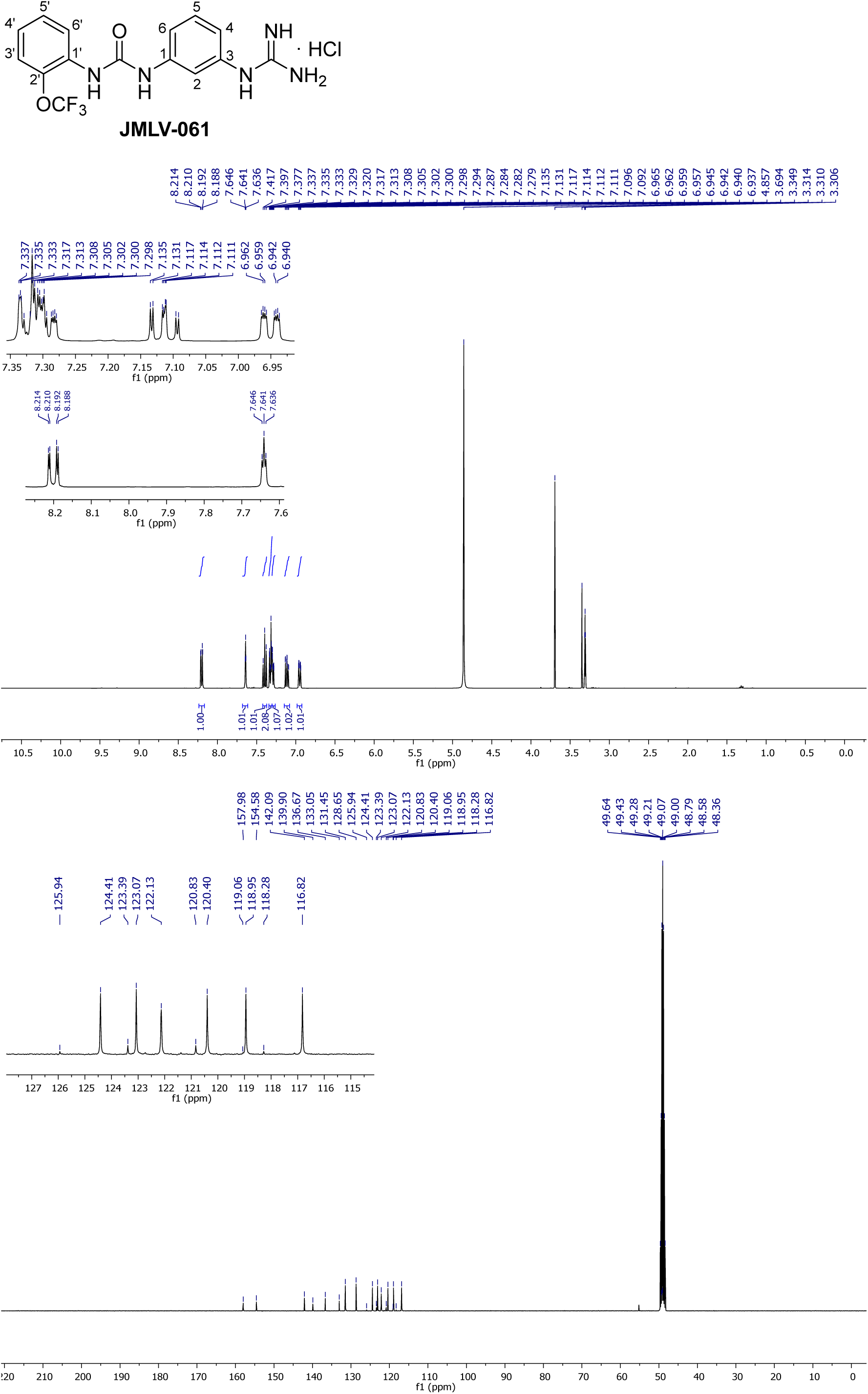

## S2 Appendix Primer and probe sequences. Oligonucleotide sequences used for sequencing or (RT-)qPCR

### S2 Appendix: Primer and probe sequences

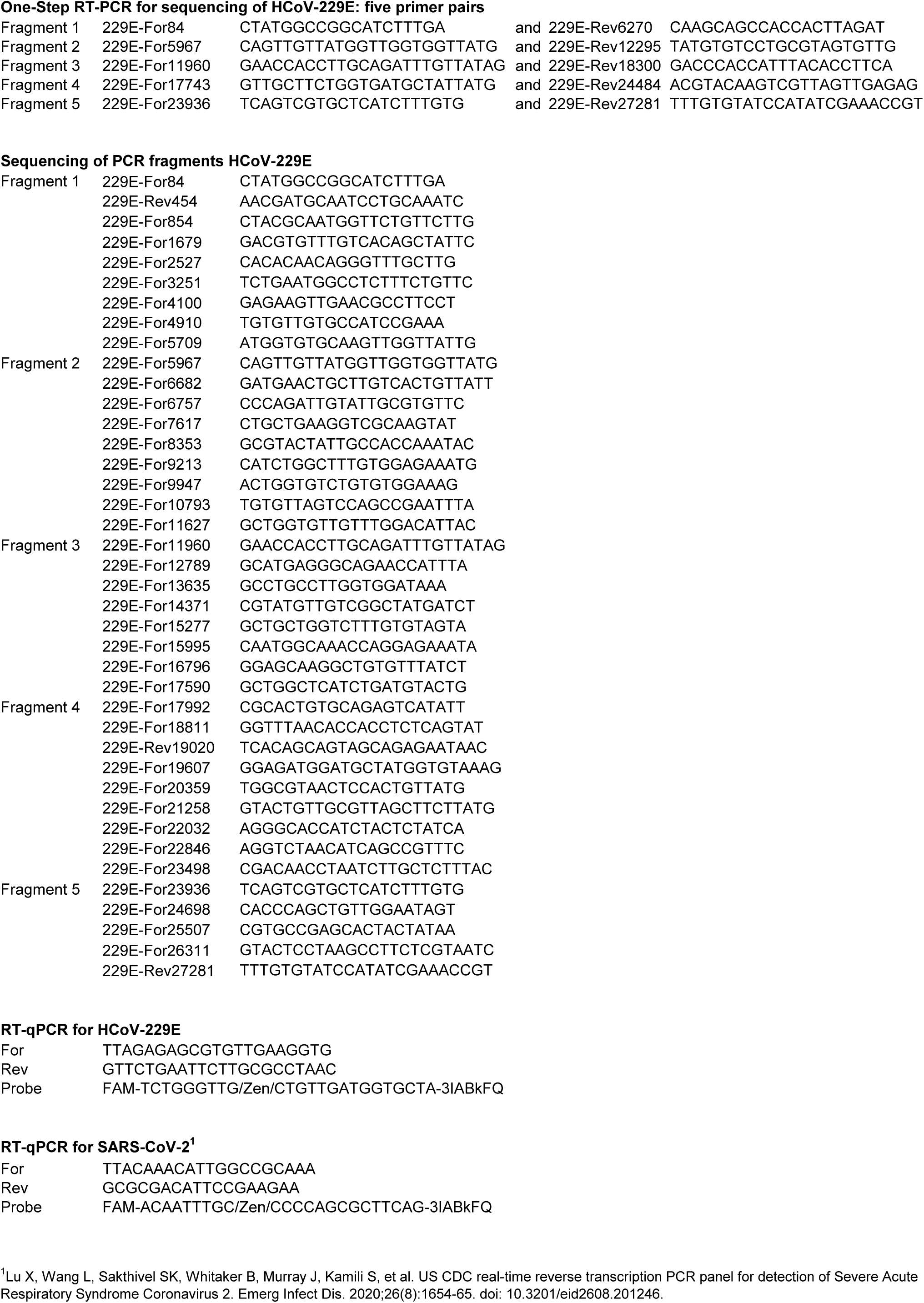

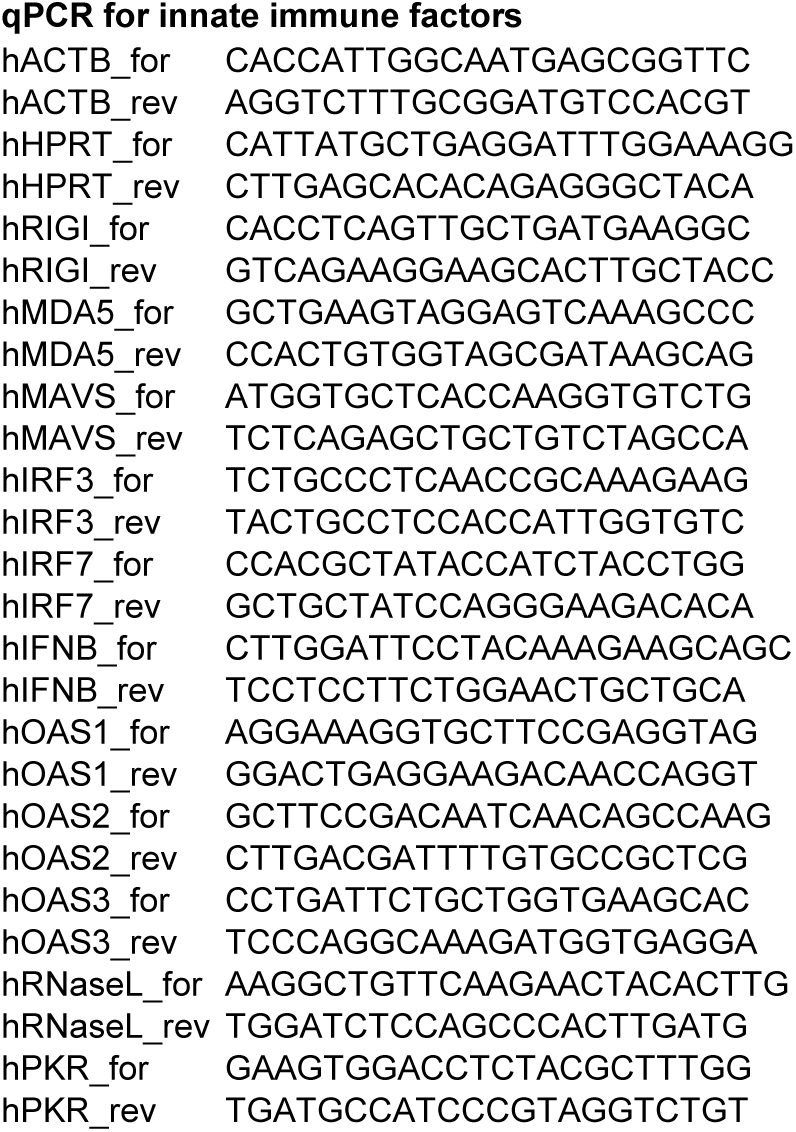

## S3 Appendix Supplementary method: automated western blot

### S3 Appendix: Protein quantification by automated western blot

To determine the intracellular protein levels of STAT1 and cleaved/uncleaved poly(ADP-ribose) polymerase 1 (PARP1), cells were lysed in RIPA lysis buffer supplemented with Halt protease inhibitor cocktail and EDTA (both from Thermo Fisher Scientific) and whole cell lysates were cleared by centrifugation. Proteins were separated by size using the 12–230 kDa Jess Separation Module (SM-W004) and bound with a primary antibody against STAT1 (Cell Signaling, catalog no. 9176, diluted 1:50) or PARP1 (Cell Signaling, catalog no. 9542, diluted 1:100). The STAT1 primary antibody was detected using the mouse detection module (DM-002, Protein Simple), while the PARP1 primary antibody was detected using the rabbit detection module (DM-001, Protein Simple). Protein separation and detection was performed according to the manufacturer’s instructions by capillary electrophoresis, antibody binding and visualization of HRP conjugates. Next, the primary and secondary antibodies were removed using the Replex Module (RP001, Protein Simple) to allow sequential total protein detection. Protein signals were visualized using Compass Simple Western software, v.6.1.0 (ProteinSimple).

**S1 Fig.**
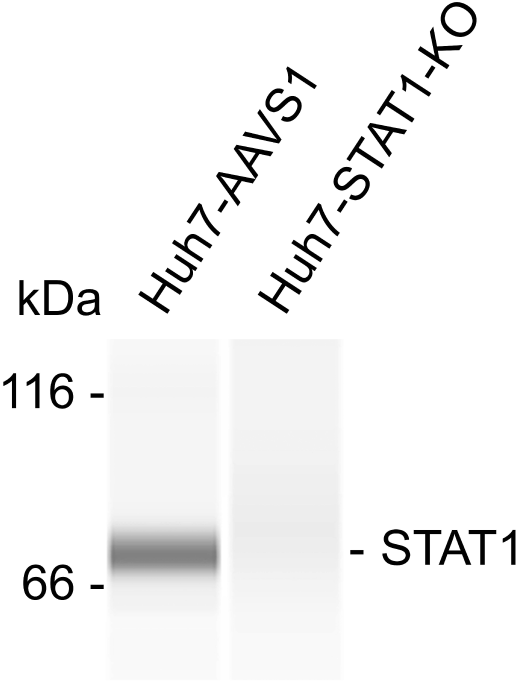
Confirmation of knock out in Huh7-STAT1-KO cells. After CRISPR/Cas9-mediated STAT1 knock out, the cells were submitted to automated western Blot. Huh7 cells expressing control sgRNAs (targeting safe harbor AAVS1) were included as STAT1-expressing control.

**S2 Fig.**
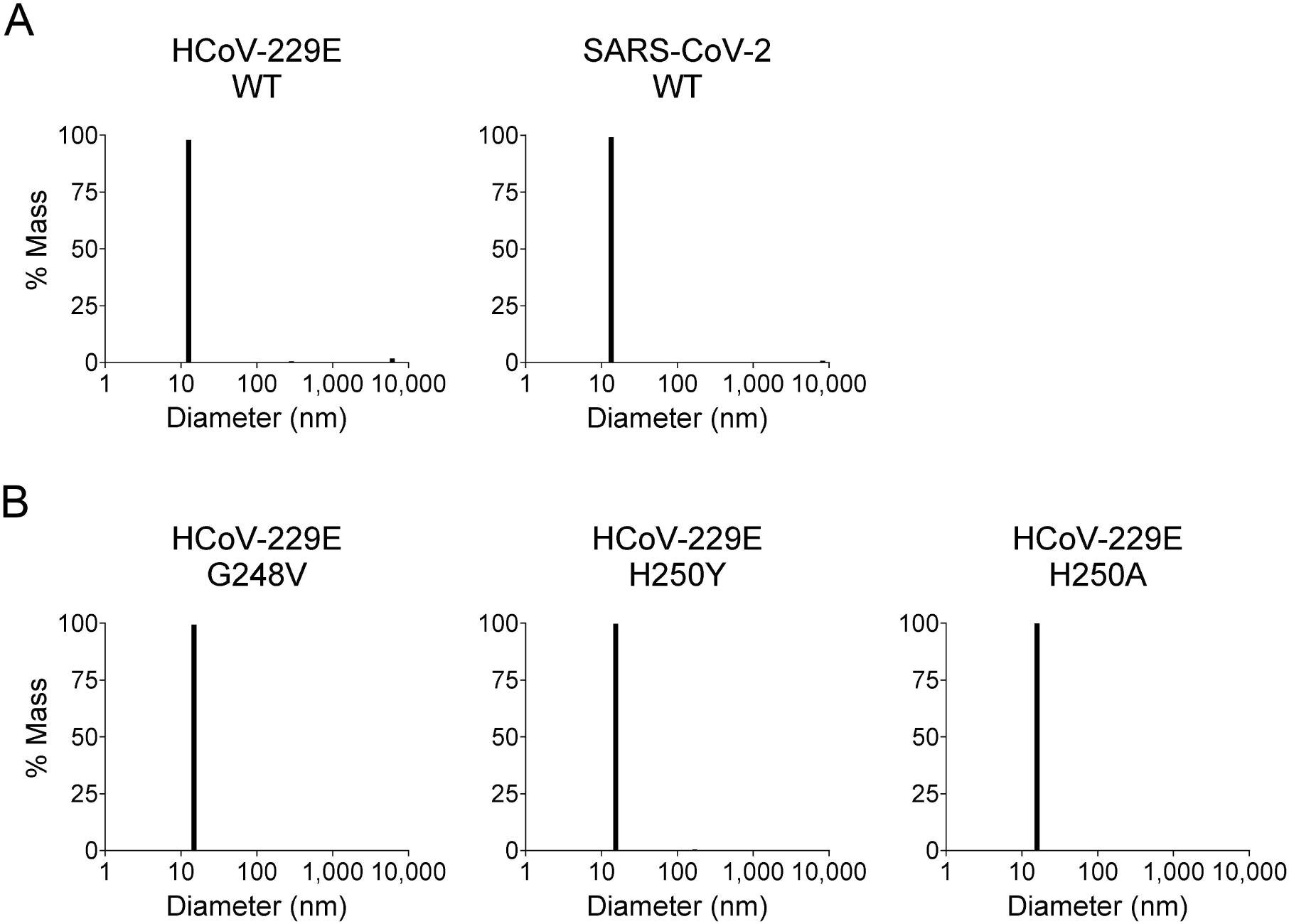
Particle size distribution of SEC-purified nsp15 samples. Particle size distribution of HCoV-229E and SARS-CoV-2 nsp15 as determined by dynamic light scattering (DLS). Cuvette-based DLS was performed on SEC-purified 0.5 mg/ml nsp15 solutions in storage buffer (25 mM HEPES at pH 8, 300 mM NaCl, 10% glycerol, and 1 mM tris(2-carboxyethyl)phosphine), using a DynaPro Nanostar and Dynamics software v7.10.1.21 (Wyatt Technologies). 15 µl of sample was mixed 5 µl water. The sample chamber was kept at 20 °C. The sample contained ≥ 98% particles with a diameter of ∼13 nm, consistent with the size of hexameric nsp15, for both HCoV-229E (polydispersity index: 9%) and SARS-CoV-2 (polydispersity index: 11.7%).

**S3 Fig.**
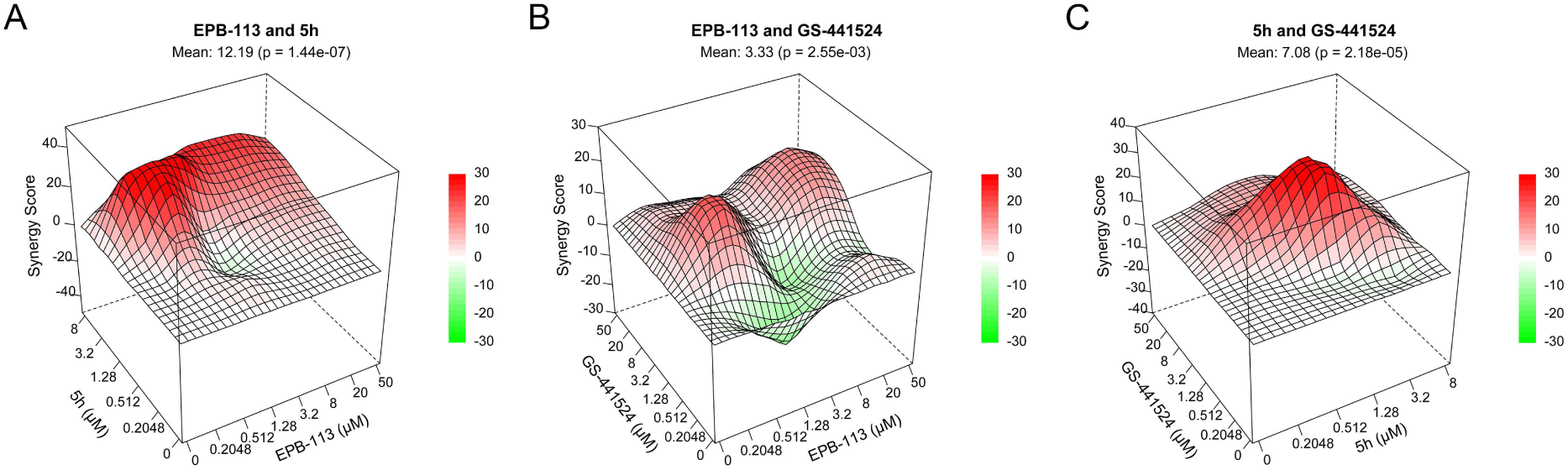
Synergism analysis of EPB-113, 5h and GS-441524. To study drug combinations, we used an alternative CPE reduction assay with HCoV-229E in HEL299 cells. An 8×8-well matrix was created, each well containing different concentrations of two compounds. Data analysis was performed with the zero-interaction potency (ZIP) model in SynergyFinder+. Mean ZIP-scores are shown on top of each graph. A ZIP-score lower than -10 indicates antagonism, between -10 and 10 corresponds with additive effects, and higher than 10 indicates synergism. The combination of EPB-113 with 5h is synergistic (A), while the combinations of EPB-113 with GS-441524 (B) and 5h with GS-441524 (C) are additive.

**S4 Fig.**
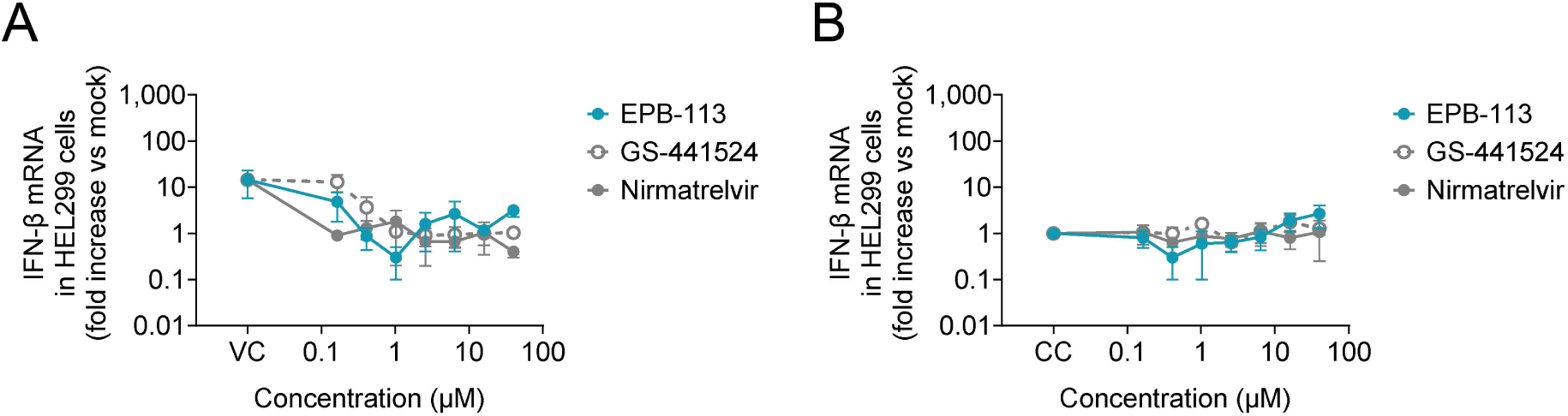
IFN-β mRNA levels upon immediate compound addition or in the absence of virus. (A) EPB-113, nirmatrelvir and GS-441524 do not increase IFN-β mRNA levels in HEL299 cells when compound and HCoV-229E virus are added simultaneously. (B) The compounds do not induce IFN-β mRNA in uninfected cells. Compound exposure time: 48 h.

**S5 Fig.**
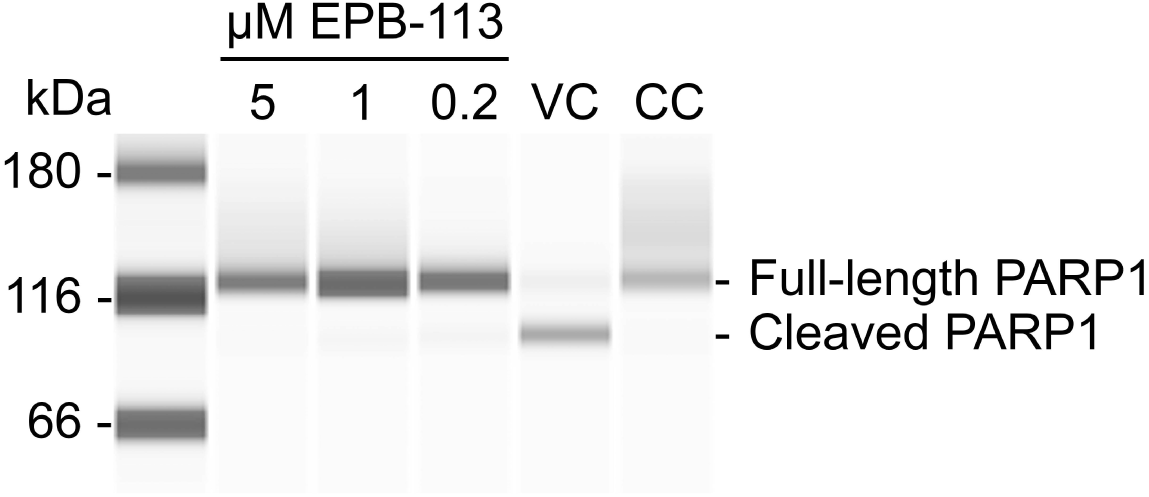
EPB-113 prevents caspase-3-mediated cleavage of PARP1 in HCoV-229E-infected HEL299 cells. Extracts were prepared at 48 h p.i. and analyzed by automated western blot (see S3 Appendix for methodology).

**S6 Fig.**
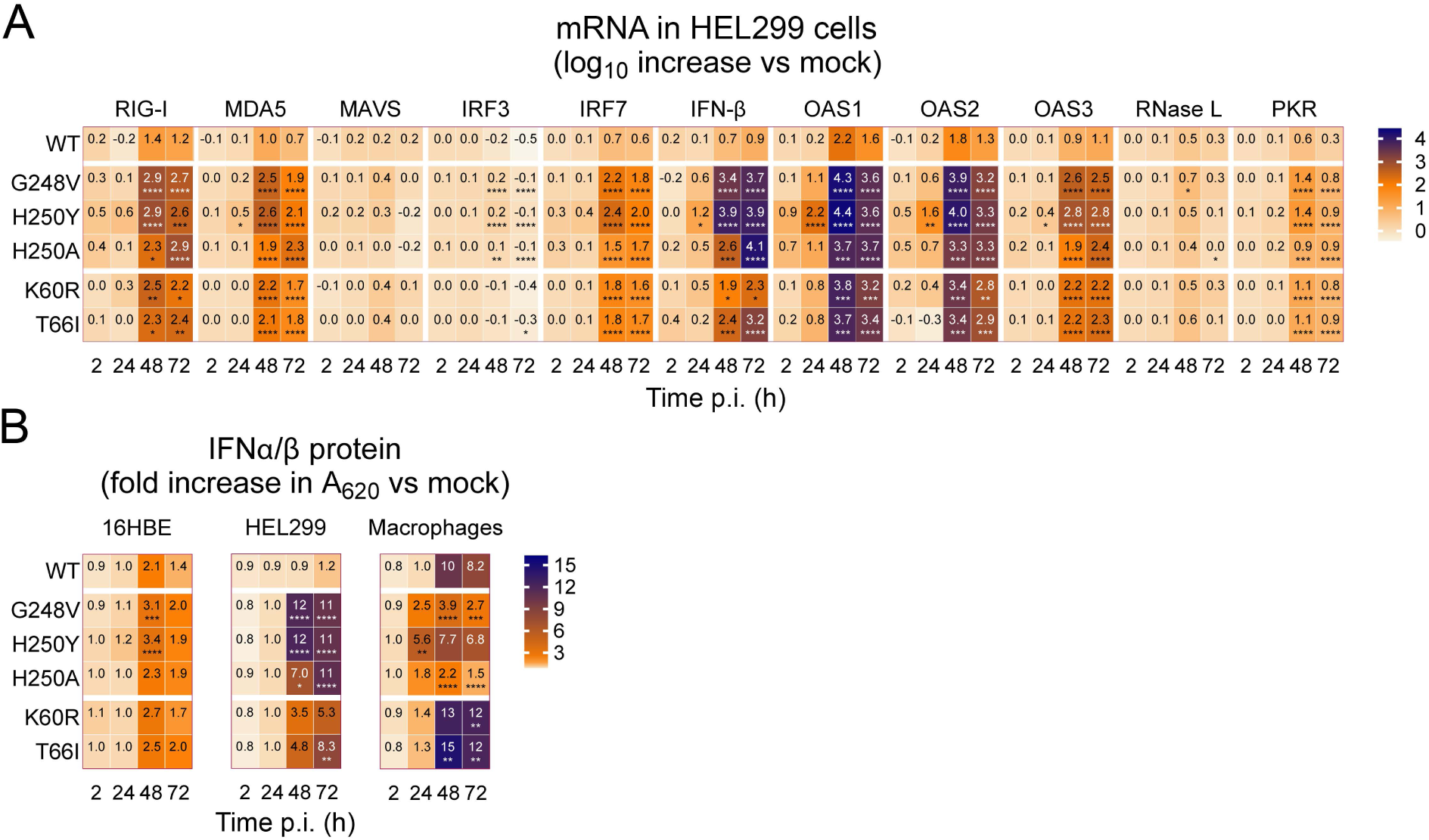
Numeric values used to create the heatmap in Fig. 9, and statistical analysis of the data.

